# Genetic architecture and evolution of color variation in American black bears

**DOI:** 10.1101/2022.08.12.503788

**Authors:** Emily E. Puckett, Isis S. Davis, Dawn C. Harper, Kazumasa Wakamatsu, Gopal Battu, Jerrold L. Belant, Dean E. Beyer, Colin Carpenter, Anthony P. Crupi, Maria Davidson, Christopher S. DePerno, Nicholas Forman, Nicholas L. Fowler, David L. Garshelis, Nicholas Gould, Kerry Gunther, Mark Haroldson, Shosuke Ito, David Kocka, Carl Lackey, Ryan Leahy, Caitlin Lee-Roney, Tania Lewis, Ashley Lutto, Kelly McGowan, Colleen Olfenbuttel, Mike Orlando, Alexander Platt, Matthew D. Pollard, Megan Ramaker, Heather Reich, Jaime L. Sajecki, Stephanie K. Sell, Jennifer Strules, Seth Thompson, Frank van Manen, Craig Whitman, Ryan Williamson, Frederic Winslow, Christopher B. Kaelin, Michael S. Marks, Gregory S. Barsh

## Abstract

Color variation is a frequent evolutionary substrate for camouflage in small mammals but the underlying genetics and evolutionary forces that drive color variation in natural populations of large mammals are mostly unexplained. The American black bear, *Ursus americanus*, exhibits a range of colors including the cinnamon morph which has a similar color to the brown bear, *U. arctos*, and is found at high frequency in the American southwest. Reflectance and chemical melanin measurements showed little distinction between *U. arctos* and cinnamon *U. americanus* individuals. We used a genome-wide association for hair color as a quantitative trait in 151 *U. americanus* individuals and identified a single major locus (*P* < 10^−13^). Additional genomic and functional studies identified a missense alteration (R153C) in *Tyrosinase-related protein 1* (*TYRP1*) that impaired protein localization and decreased pigment production. Population genetic analyses and demographic modeling indicated that the R153C variant arose 9.36kya in a southwestern population where it likely provided a selective advantage, spreading both northwards and eastwards by gene flow. A different *TYRP1* allele, R114C, contributes to the characteristic brown color of *U. arctos*, but is not fixed across the range.

**HIGHLIGHTS:**

- The cinnamon morph of American black bears and brown bears have different missense mutations in *TYRP1* that account for their similar coloration
- *TYRP1* variants in American black bears and brown bears are loss-of-function alleles associated with impaired protein localization to melanosomes
- In American black bears, the variant causing the cinnamon morph arose 9,360 years ago in the western lineage where it provides an adaptive advantage, and has spread northwards and eastwards by migration

## INTRODUCTION

Variation in animal color is a longstanding system with which to investigate gene action and evolution. In mammals, nearly all pigment is produced by melanocytes via polymerization of dihydroxyindole derivatives to yield insoluble melanins that are transferred to overlying keratinocytes in skin and/or hair. Genetic variation in melanin biosynthesis or transfer gives rise to hair, eye, and skin color differences, and can underlie positive selection [1, 2]. Mutations of melanin biosynthesis underlie a number of conditions associated with impaired fitness or a disease (e.g. human albinism), or unusual color morphs of large mammals that are specifically targeted for trophy hunting [3].

Identifying and understanding the genetic causes and consequences of mammalian color variation is of special interest in large mammals for two reasons. First, by contrast to small mammals in which color morphs are often selected as a means of predator avoidance, the evolutionary forces underlying color variation in large mammals is less apparent and occasionally controversial. Second, there are often charismatic and/or cultural values associated with certain species and color morphs such as the black jaguar and white tiger. Despite their common name, American black bears (*Ursus americanus*) occur in black, cinnamon, brown, blond, grey, and white morphs; similarly, brown bears (*U. arctos*) occur in blond, brown, dark chocolate, white, and ruddy. The cinnamon morph of *U. americanus* has a rich history in popular culture [4] and systematics, and has been suggested to have adaptive value, either by aiding in thermoregulation in hotter and drier climates of the southwest, or by mimicry of *U. arctos* in areas where the two species are sympatric. Cinnamon has a distinct latitudinal gradient in western North America, with a high frequency in the southwest that declines moving northwards, and is at low frequency in the east [5]. This phenotypic variation is consistent with genomic analyses that indicate the deepest phylogeographic structuring occurs between the eastern and western lineages across North America [6, 7]. Further, admixture between the lineages has been identified across the northern range and suggested to be a product of post-glacial range expansion, and east-west gene flow across contiguous habitat in modern Canada [7].

The distinct spatial distribution of cinnamon *U. americanus* presents three alternative phylogeographic hypotheses. First, the causative mutation arose in the western lineage after the split from the eastern lineage. Second, the variant predates lineage divergence, but was lost during eastward range expansion. Third, introgression of the causative variant from *U. arctos* into *U. americanus* occurred in a western lineage population.

We used a genome-wide approach and identified the causative variant that produces cinnamon morphs in *U. americanus*; then applied molecular and population genetic approaches to explore its functional impact and demographic history. We identified a missense alteration in *Tyrosinase-related protein 1* (*TYRP1*) and show that it interferes with melanogenesis and melanin synthesis. We find the *U. americanus TYRP1* variant was not introgressed from *U. arctos*; instead, a different *TYRP1* variant contributes to their characteristic pigmentation phenotype. Remarkably, the variant responsible for cinnamon *U. americanus* is identical to one previously described as a cause of oculocutaneous albinism (OCA3) in humans, often associated with nystagmus and reduced visual acuity.

Our analyses suggest that the *U. americanus TYRP1* variant arose within the western lineage of *U. americanus* and likely provided a selective advantage in the southwest. These results illustrate how Mendelian variation in melanogenesis can underlie iconic phenotypes and inform our understanding of color variation and recent evolution in large carnivores.

## RESULTS and DISCUSSION

### Quantitative and Chemical Characterization of Bear Hair Color

In addition to cinnamon, light-colored morphs of *U. americanus* have been described as blond, light brown, or chocolate; thus, the results presented reflect all of these color morphs. We measured reflectance of hair samples from 391 *U. americanus* and 33 *U. arctos* from across North America. Most *U. americanus* qualitatively described as black based on photographs exhibited reflectance values < 50 (median = 5), whereas animals scored as cinnamon had broadly ranging values (median = 67; Figure 1A); overall, the distribution of reflectance varied continuously from 0.5–182 (possible range: 0–255; Figures 1A, S1). Reflectance in *U. arctos* also varied continuously (median = 48). Thus, we consider the reflectance measure a quantitative variable.

**Figure 1-.**
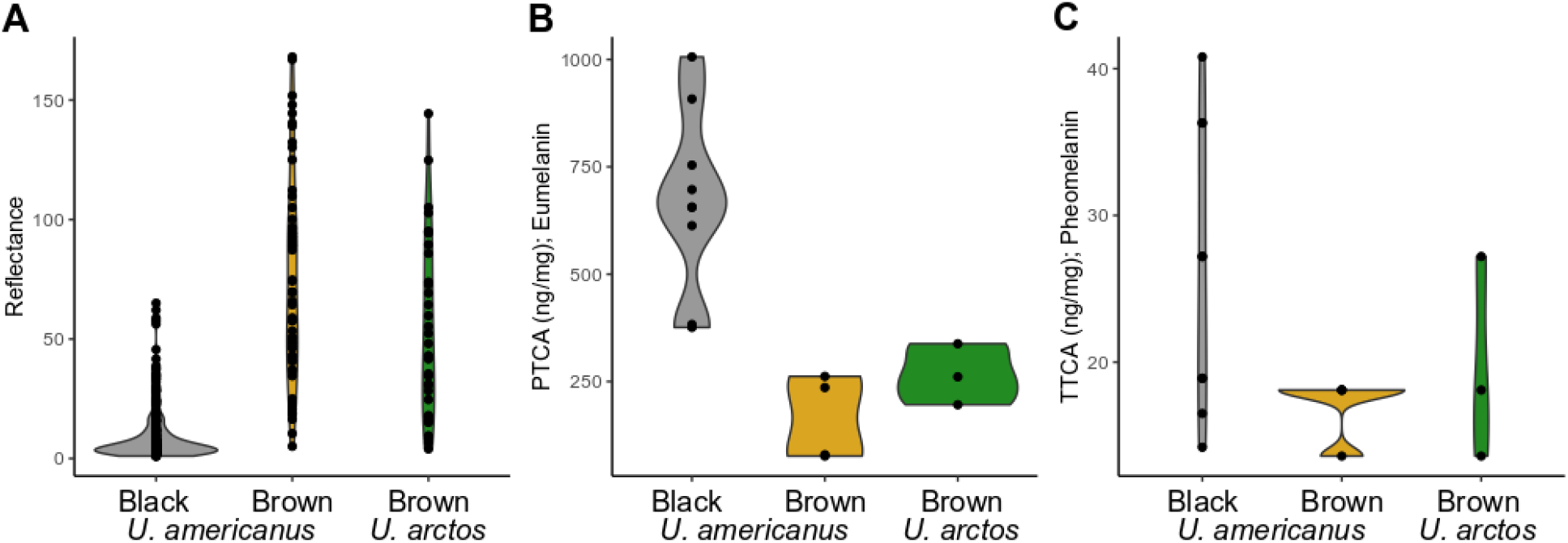
Characterization of bear hair color. (A) Violin plots of hair reflectance for two bear species (*Ursus americanus* [n = 327] and *U. arctos* [n = 33]) categorized by qualitative phenotyping from photos as either black or brown animals (black *U. americanus*- grey; brown *U. americanus*- yellow; brown *U. arctos*- green). Chemical analysis via alkaline hydrogen peroxide oxidation followed by HPLC of 13 *U. americanus* and three *U. arctos* individuals’ hair for the concentration of (B) eumelanin (as PTCA; ANOVA *F*=14.512; *p*<0.001) or (C) pheomelanin (as TTCA; ANOVA *F*=2.0297; *p*=0.17). PTCA concentration in hair ranged from 75–1010 ng mg^-1^, where TTCA was limited to 13–40 ng mg^-1^ in *U. americanus*. Specifically, two-tailed t-tests between *U. americanus* black and brown animals (*p*<0.001), and *U. americanus* black and *U. arctos* were significantly different (*p*<0.001); whereas, no difference was observed between species with brown coloration (*p*=0.17). In *U. arctos*, PTCA and TTCA ranged respectively from 200–340 ng mg^-1^ and 13–27 ng mg^-1^. Both species indicate a dilution of eumelanin, and not an increase in pheomelanin, produces the characteristic hair-lightening.

We determined if hair from cinnamon *U. americanus* and *U. arctos* was caused by dilution of eumelanin or increased production of pheomelanin using HPLC [8, 9] of the different pigment types in awn hair samples. As depicted in Figures 1B-C, the major determinant of light-colored hair is reduced amounts of eumelanin in both bear species.

### Allelic Identification of TYRP1 Variants

We sequenced the genomes of 24 *U. americanus* to ≥30x depth and an additional 166 animals to 1.3x depth (Table S1), mapped to the reference genome [10], then imputed missing loci in the low coverage samples using the high coverage individuals as a reference panel [11]. A genome-wide association study (GWAS) [12] for quantitative hair reflectance values from 151 bears identified 120 single nucleotide variants (SNVs) above a significance threshold of 10^−8^ (Figure 2A, Table S2). A single peak had the lowest *P*-values and spanned the pigmentation gene *TYRP1* (Figure 2B), in which we identified a significant (*P*=1.99 × 10^−13^) G to A SNV, HiC_scaffold_24:6724152g>a that predicts a missense alteration, p.Arg153Cys, referred to in what follows as *TYRP1*^*R153C*^. Nine additional SNVs had lower *P*-values and were 130kb downstream from the *TYRP1* transcription start site (Figure 2B); however, after accounting for *TYRP1*^*R153C*^ genotypes as a covariate, the significance of these sites fell below the genome-wide threshold (Figure S2). Further, haplotype analysis [13] identified low diversity in the Nevada population, thus we used these samples as input into HAPLOVIEW [14] and identified a single 97kb haploblock carrying *TYRP1*^*R153C*^ (Figures 2B, S3). Recombination is apparent in populations outside of the southwest, suggesting that *TYRP1*^*R153C*^ is derived from a single mutational event.

**Figure 2-.**
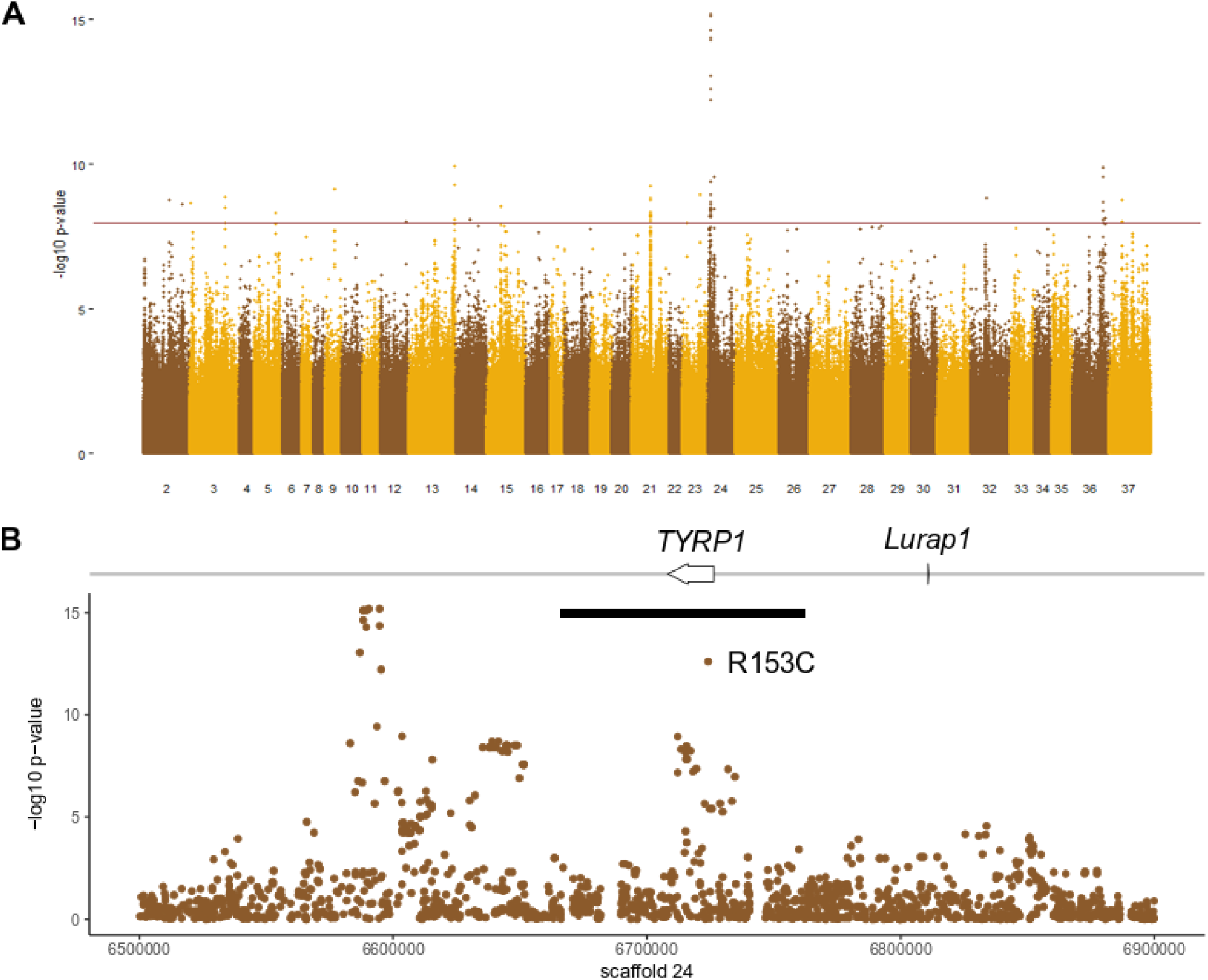
GWAS of both high and low coverage WGS data from *Ursus americanus* (n = 151) to identify loci associated with coat color. (A) A genome-wide Manhattan plot with a significance cut-off of 10^−8^ (horizontal brown line) identified a single strong peak on scaffold 24 (additional peaks in Table S6). (B) A detailed view of scaffold 24 surrounding the peak identified two genes, including *TYRP1*. The black bar denotes the length of the haplotype identified within the Nevada population that contains the R153C derived allele.

We explored coding sequence variation in *TYRP1* and 12 additional pigmentation genes in *U. americanus, U. arctos*, and *U. maritimus* (polar bear; Table S3). For *U. americanus*, we identified 46 missense or nonsense polymorphisms within 11 genes (Table S4); among these, only the *TYRP1*^*R153C*^ variant was correlated with color (Figure S4A) and predicted to be deleterious. We identified 19 missense variants between *U. americanus* and *U. arctos* (Table S5), and 33 missense or nonsense variants between *U. arctos* and *U. maritimus* (Table S6). The *U. americanus* R153C variant was not observed in *U. arctos* or *U. maritimus*; however, a different *TYRP1* variant was identified in some *U. arctos*, HiC_scaffold_24:6725331g>a, that predicts a p.Arg114Cys substitution (*TYRP1*^*R114C*^). Like *TYRP1*^*R153C*^ in *U. americanus, TYRP1*^*R114C*^ in *U. arctos* was predicted to be deleterious (Table S5; Figure S4) and, as shown below, interferes with TYRP1 function.

We also examined *TYRP1* variation at orthologous positions in other species. In humans, loss-of-function for *TYRP1* causes a rare form of albinism, rufous oculocutaneous albinism (OCA3), observed mostly in individuals of African or Puerto Rican ancestry, and characterized by reddish skin and hair and frequent visual abnormalities [15]. Among 15 individuals with clinical albinism and mutations of *TYRP1*, R114C and R153C were each observed once as compound heterozygotes [16]. The R153C variant is found at a relatively high frequency in individuals of European ancestry, 7.91×10^−4^ but is classified as a variant of uncertain significance [17, 18]. In laboratory rats the R114C allele is likely responsible for a spontaneous brown mutation that is fixed in the Brown Norway strain [19].

### Functional Characterization

We investigated functional consequences of the *TYRP1* R153C and R114C variants by measuring melanogenesis and TYRP1 localization in mouse melan-b cells, which are derived from *TYRP1*^*b/b*^ mutant mice. We stably expressed in these cells human or mouse TYRP1 with or without the R153C or R114C variants, or mCherry-syntaxin 13 (mCh-STX13) as a negative control, from recombinant retroviruses. Using a colorimetric melanin content assay (Figure 3A), wild-type (WT) human TYRP1 rescued pigmentation relative to untransfected (-) melan-b cells to nearly the same level as WT (melan-Ink4a) melanocytes. By contrast, the R153C and R114C variants rescued pigmentation in melan-b cells respectively only 52.0% and 14.1% as well as WT human TYRP1, and 46.0% and 6.8% as well as WT mouse TYRP1. This is consistent with melanin quantification in bear hair detected by HPLC (Figure 1B).

**Figure 3-.**
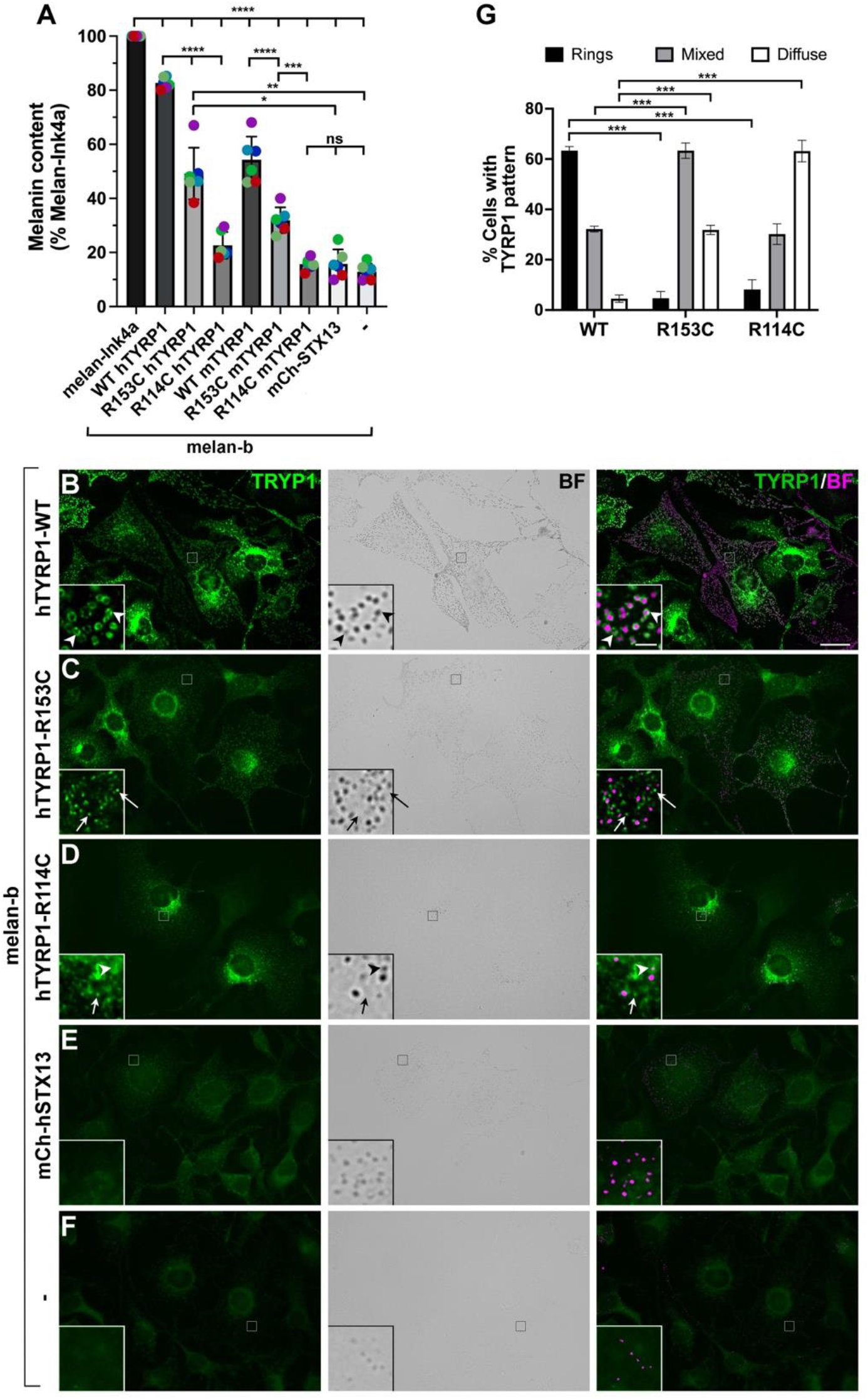
(A) TYRP1-deficient melan-b cells that were untransduced (-) or stably expressing mCh-STX13 or the WT, R153C, or R114C variants of human (h) or mouse (m) TYRP1, were analyzed by quantitative melanin content assay and normalized to protein content. Data are presented as a percent normalized melanin content relative to that of WT melan-Ink4a cells from six experiments (colored dots), each performed in duplicate, and analyzed by two-way ANOVA and Tukey’s multiple comparison test. Indicated untransduced (-; F) or stable melan-b transductants (B-E) were analyzed by immunofluorescence microscopy for TYRP1 (left, green) and bright field microscopy (BF) for pigment granules (middle); right, overlay with pigment granules pseudocolored magenta. Insets, 7X magnification of boxed regions (intensities of TYRP1 and pigment granules optimized to better visualized overlap). Arrowheads, TYRP1 in rings around pigment granules; arrows, TYRP1 in separate punctate structures. (G) Quantification of cellular pattern of TYRP1 as predominantly rings, diffuse/punctate, or mixed among transduced cells expressing each of the TYRP1 variants. Data are from four experiments analyzing 150 of each cell type per experiment, and were analyzed by two-way ANOVA with Dunnett’s multiple comparison test relative to WT TYRP1.

To test if the defect in pigmentation reflected impaired localization of the variants to melanosomes, we visualized the distribution of the human TYRP1 variants within melan-b cells by immunofluorescence and bright field microscopy (Figure 3B-F). Only background labeling for TYRP1 was detected in untransduced melan-b cells (-; Figure 3F) or cells transduced with mCh-STX13 (WT, Figure 3E). Wildtype human TYRP1 was detected predominantly in “rings” around pigment granules detected by bright field microscopy (Figure 3B,E,G; see arrowheads; G). By contrast, the R114C variant was primarily detected in punctate and diffuse structures that did not overlap the lighter pigment granules in these cells (Figure 3D, arrows; 3G). The R153C variant was mostly detected in a mixture of ring-like and diffuse/ punctate patterns, (Figure 3C,G). Both variants were also generally expressed at lower levels than WT TYRP1. Together, these data indicate that *TYRP1*^*R153C*^ and *TYRP1*^*R114C*^ are hypomorphs. Because the TYRP1 lumenal domain is required for proper melanosome localization [20], a likely pathophysiologic mechanism to explain eumelanin dilution in bears is that substitution of Cys for Arg at residues 114 or 153 leads to protein misfolding, resulting in enhanced protein degradation and impaired trafficking to and/or retention within melanosomes.

### Gene Action and Spatial Distribution of TYRP1^R153C^

For *U. americanus*, examining hair reflectance as a function of *TYRP1*^*R153C*^ genotype in 317 individuals revealed semidominant gene action, with a median reflectance value for heterozygous G/A (Arg/Cys) individuals, 50.3, that was intermediate between values for homozygous G/G (Arg/Arg) and A/A (Cys/Cys) individuals of 4.7 and 94.0, respectively (inset, Figure 4).

**Figure 4-.**
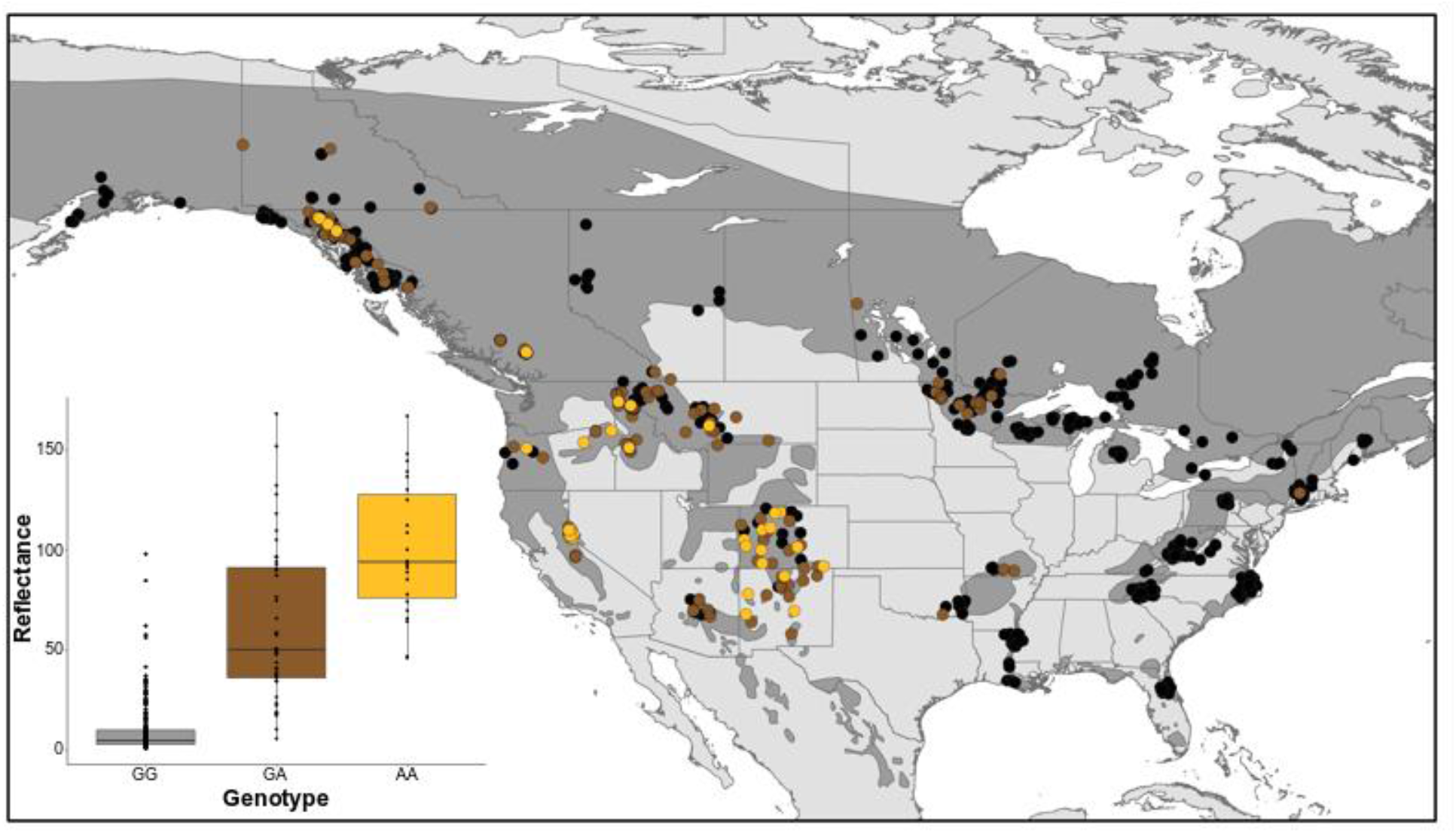
Quantitative and spatial assessment of *TYRP1*^*R153C*^ in *Ursus americanus*. (Inset) Hair color reflectance was compared to the SNP genotype (n = 317). We used t-tests to compare reflectance values between genotypes: homozygous ancestral and heterozygous *P*=2.02 × 10^−11^; and heterozygous and homozygous derived *P*=1.41 × 10^−4^. These results indicate the allele acts with semidominance. The geospatial pattern of R153C across the range (n = 906) where color denotes genotype (black: homozygous ancestral GG; brown: heterozygous GA; gold: homozygous derived AA), with the species range shown in dark grey.

We determined the spatial distribution of *TYRP1*^*R153C*^ in 906 samples across the species’ geographic range. A high frequency of the derived allele was apparent in the southwest, declining northward along the Rocky Mountains into Southeast Alaska and Yukon Territory (Figure 4). By contrast, a low frequency of the derived allele was apparent in the eastern lineage, primarily limited to the Great Lakes region and a single sample in Connecticut, USA. The presence of the derived allele in Missouri, Arkansas, and Oklahoma, USA was concordant with the known translocation history of bears from Minnesota into Arkansas in the 1960s [21] and the high proportion of genetic diversity from the Great Lakes region now found in the Central Interior Highlands [22]. Spatial distribution of *TYRP1*^*R153C*^ was consistent with the phenotypic distribution of non-black *U. americanus* as estimated from a survey of 40,000 animals in the 1980s [5]. Notably, *TYRP1*^*R153C*^ was associated with bears qualitatively described as cinnamon, chocolate, or light brown, and therefore accounts for the majority of color diversity among *U. americanus*.

### Demography

Our *TYRP1* haplotype tree (Figure S4A) indicated that introgression of *U. arctos* alleles into *U. americanus* did not provide the genetic material for light coloration in the latter species. We investigated the relationship of the *TYRP1*^*R153C*^ mutation to *U. americanus* demographic history. First, we used *runtc* [23] to estimate the first coalescent (a proxy for allele age) of the variant along the scaffold and obtained a time of 9.36kya. Second, we ran MSMC-IM [24] from whole genome resequencing (WGS) of six individuals which estimated that divergence between the western and eastern lineages began 100kya with cessation of gene flow 22kya (Figure 5A,B).

**Figure 5-.**
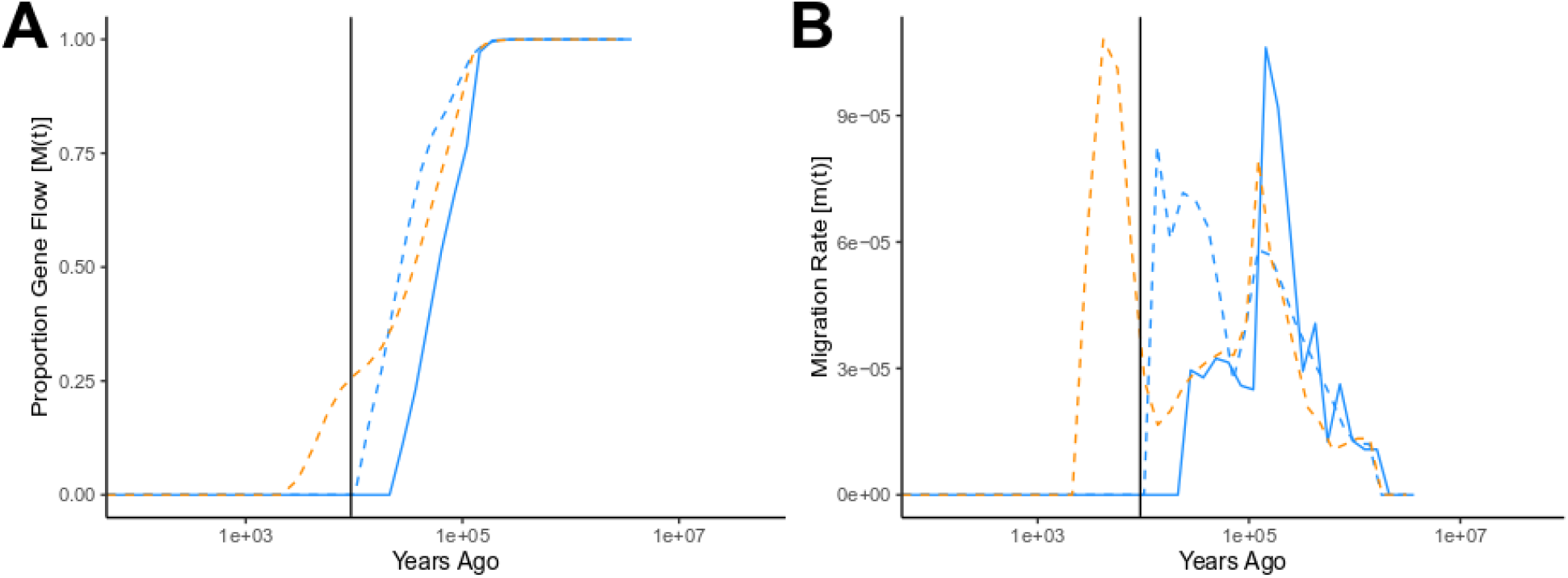
*Ursus americanus* demographic estimates over time. (A) Approximate completion of divergence and (B) bidirectional migration between lineage pairs including: eastern and western (solid blue line), eastern and Southeast Alaska (dashed blue line), and western and Southeast Alaska (dashed orange line). The vertical black line represents the estimated allele age of *TYRP1*^R153C^ (also see Figure S5).

Taken together with additional coalescent and range expansion analyses (Figures S5, S6), our results suggest that R153C arose in a western lineage population of *U. americanus* and spread via gene flow across the range, including contemporary eastward movement too young to be captured using these coalescent estimates.

### Selection Analyses

We used population genetic modeling and simulations to investigate whether clinal variation of R153C allele frequency was more likely to be explained by selection or genetic drift. We compared models with a range of dominance (*h*) and selection (*s*) coefficients (Figure S7). As depicted in Figure 6, the clinal pattern of allele frequencies is consistent with weak selection (0.005 < *s* < 0.01) and is unlikely to be explained by genetic drift (*s* = 0). Despite the long haplotype carrying the derived *TYRP1*^*R153C*^ allele, there is no statistical evidence for a selective sweep (Figure S8-S10), likely due to the combination of weak selection and low effective population size within the southwest population. In humans, *SLC24A5* drives the main axis of geographic color variation in Caucasians and has a large *s* of 0.16 [25], although other genes, *TYRP1* among them, have weaker coefficients of 0.01-0.05 [25, 26].

**Figure 6-.**
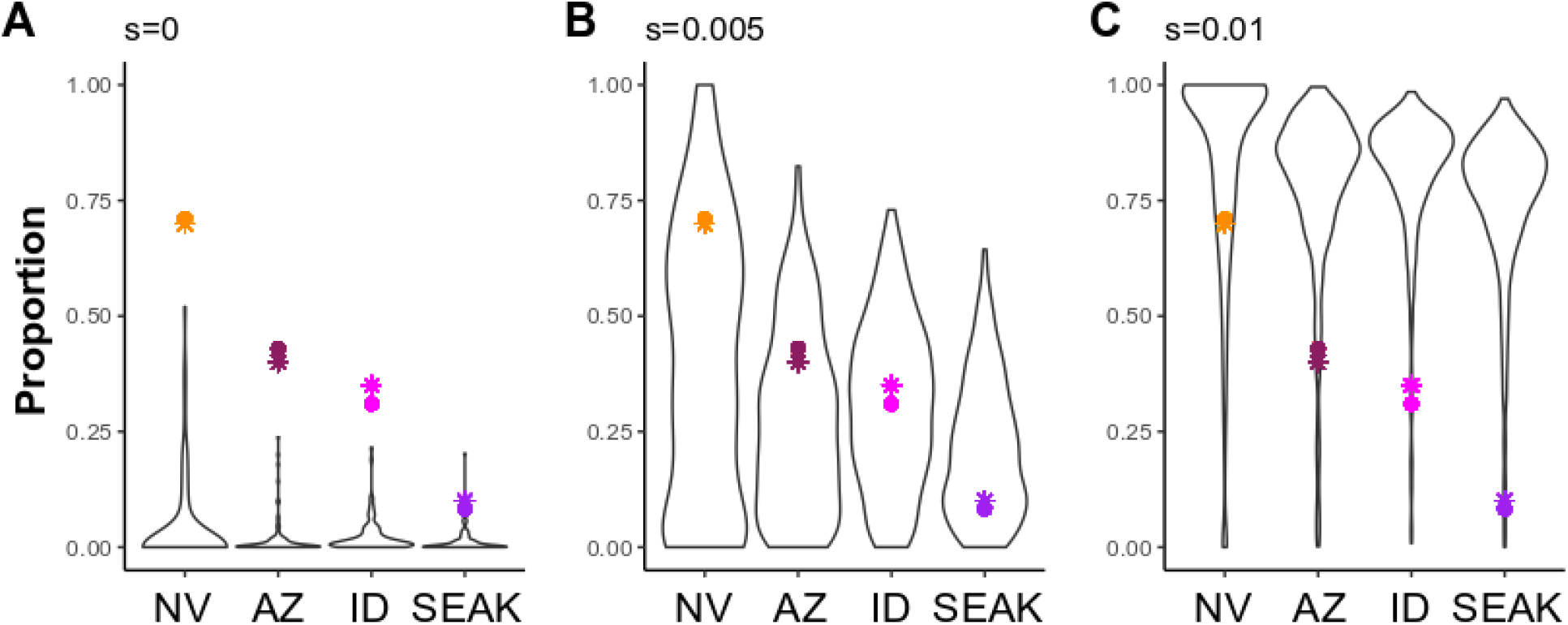
Simulation models for the frequency of a derived allele (*h* = 0.75) following 1,440 generations in four populations (Nevada: NV; Arizona: AZ; Idaho: ID; and Southeast Alaska: SEAK) connected via gene flow (see Figure S11). Models varied by selection coefficients (*s*; A- 0; B- 0.005; and C- 0.01), and accounted for population expansion and bottlenecks. Each simulation model was run for 1,000 iterations and panels represent runs in which the derived allele did not go extinct within a run; thus, sample size varies (A- 95; B- 394; and C- 614; see Figure S7-bottom row). Colored points represent R153C genotypic frequencies (circles) estimated in this study for each respective population, or phenotypic frequencies (stars) inferred from contour maps presented in Rounds (1987).

Two hypotheses about the selective forces acting upon the cinnamon morph of *U. americanus* have been proposed. First, lighter coloration aids in thermoregulation of bears in the hotter and/or drier climates of the southwest; second, the cinnamon morph is mimetic with *U. arctos* where these species are sympatric. We used a Bayesian model for correlation of allele frequencies and environmental factors, BayEnv2 [27], to test for a relationship between R153C allele frequency and climate, geography, or the presence/absence of *U. arctos* prior to anthropogenic extirpation, but identified no notable Bayes Factors (Table S7). Although this approach cannot rigorously exclude thermoregulation or mimicry as a force driving selection, it is possible that an additional, as yet, untested mechanism underlies weak selection such as crypsis in open canopy environments. For example, individuals color matched within their environment has been suggested to decrease infanticide and/or predation in the Giant Panda [28].

### Implications for Infraspecific Taxonomy in American Black Bears

Historically brown coat color has been used as a defining phenotype for delimiting four of the 16 *U. americanus* subspecies: *U. a. amblyceps* (southwest/southern Rocky Mountains), *U. a. cinnamomum* (northern Rocky Mountains), *U. a. luteolus* (Louisiana), and *U. a. machetes* (western Mexico) [29-31]. Further, the original description of *U. luteolus* from 1821 [30] notes the cinnamon morph ranging from Virginia to Louisiana, where it is not presently found. Our results indicate that the geographic distribution of brown coats is being driven by recent and ongoing gene flow, and suggests that cinnamon morphs will increase in frequency within the eastern lineage in the future. We suggest the young age of the causative variant and rapid spread via gene flow make this a poor character trait for infraspecific delimitation.

### Conclusion

Here we show that the cinnamon morph of the American black bear shares phenotypic similarity with brown bears in their coat coloration, and demonstrate that eumelanin dilution causes this similarity. We identified two independently evolved and species-specific alleles within *TYRP1* that result in arginine to cysteine residue changes that disrupt protein localization to the melanosome. Our functional assay shows that the *U. arctos* R114C change has a greater effect on pigment production in melan-b cells than the *U. americanus* R153C; yet R114C has not fixed within the species (Figure S4B) suggesting it may modify color without being the initial causative locus. In contrast, the *U. americanus* R153C is responsible for most color variation within the species. Neither variant appears detrimental to fitness in bears unlike in humans with OCA3; instead R153C appears to be under weak positive selection. Crypsis as an adaptive mechanism has generally explained why prey species and ambush predators color match within their environments; here, we suggest crypsis as a broader adaptive mechanism for large bodied species.

## METHODS

### Sample Collection

We partnered with state, provincial, and federal wildlife agencies, and university partners in North America which collected hair, tissue (i.e., muscle or blood), and photos from bears that were hunter harvested, vehicle killed, or live animals captured for other research studies or management purposes. Not all samples contained all three sampling elements (hair, tissue, and photos). Photographs of bears confirmed all samples were adults; cubs may turn from brown to black following a molt, although black to brown transitions have not been observed (personal observation DL Garshelis). Samples were collected from 2015–2020 and georeferenced. The qPCR analysis (see below) incorporated previously collected bear samples [32] that were also georeferenced but without phenotypic data.

### Hair Analyses

We mounted one awn hair from 391 individuals onto slides, then captured an image from the middle of the sample under 20x magnification using an Olympus BX63 microscope at the Integrated Microscopy Center at the University of Memphis. We viewed all samples with transmitted light brightfield illumination, 488.3 µs exposure time, and 185 LED brightness. We analyzed images in ImageJ [33] using the RGBMeasure function on a 46 × 46 pixel section of the hair cortex, and recorded the mean for the red, green, and blue color channels.

We quantified the amount of eumelanin and pheomelanin in 13 *U. americanus* and three *U. arctos* individuals. Samples were selected based on both geographic location and visual inspection of hair color under a compound microscope to select for observed variation (*U. americanus*: black = 9; brown = 4; *U. arctos*: dark brown = 2, brown = 1). We followed the melanin determination protocols [8, 9]. Eumelanin and pheomelanin concentrations are reported as the concentrations of PTCA (pyrrole-2,3,5-tricarboxylic acid) and TTCA (thiazole-2,4,5-tricarboxylic acid) which are, respectively, the specific degradation products for each melanin.

### Whole Genome Sequencing and Variant Calling

#### High Coverage

We sequenced 15 *U. americanus* and six *U. arctos* individuals to ∼30x depth (Table S1). Specifically, 350bp insert libraries were prepared using the NEB Next Ultra II DNA kit, then sequenced (150bp paired-end) on an Illumina NextGen by Novogene (Chula Vista, CA). We sequenced nine additional *U. americanus* individuals to 50x depth using Illumina HiSeqX at HudsonAlpha (Huntsville, AL). We also downloaded four *U. arctos* and two *U. maritimus* WGS sequences from the NCBI SRA to use in downstream analyses (Table S1). For all species, we mapped reads to the *U. americanus* reference genome [10] using BWA-MEM v0.7.17 with default parameters. We marked duplicates, then called variants for each sample using the GenotypeGVCFs function in GATK v4.1.8.0 where we set the heterozygosity parameter to 5.0×10^−4^. For each species, we combined the samples using CombineGVCFs before joint genotyping using GenotypeGVCFs in GATK. We filtered the dataset using BCFTOOLS v1.9 by requiring minimum and maximum depth (-i ‘MIN (FMT/DP) > 4 & MAX (FMT/DP) < 300’), minimum quality score (-i FMT/GQ >= 30), and biallelic SNPs (-m 2 -M 2 -v snps). The total number of SNPs for *U. americanus* was 8,885,511. We phased the data from each species separately using BEAGLE v5.1 [34, 35].

#### Low Coverage

We sequenced 166 *U. americanus* animals to an average of 1.3x depth. Specifically, we followed the manufacturer’s instructions for the Illumina DNA Prep Kit except that we diluted the reactions to 20% volume and started with 20 ng of genomic DNA. Two pooled libraries were constructed and each was sequenced on one lane of an Illumina 4000 at Novogene (Davis, CA). We mapped reads to the *U. americanus* reference and called variants as described above. The draft *U. americanus* (2n = 74) genome was assembled into 94,016 scaffolds; we analyzed the longest 36 scaffolds (and removed the X chromosome, HiC_scaffold_1). We used our high coverage WGS samples as a reference panel for joint imputation and phasing of the low coverage data using GLIMPSE [11].

### GWAS and Haplotype Characterization

To identify loci associated with the brown phenotype, we ran GEMMA v0.98.1 [12]. Specifically, we combined our high and imputed low coverage datasets, then used the quantitative hair reflectance values (green channel of RGB measurement) as the phenotype (n = 151). We ran a PCA on the relatedness matrix created within GEMMA, and used the first two PC axes as covariates to account for population structure in the analysis. Our GWAS resulted in 23 genomic regions with *P*-values lower than 10^−8^, although a single peak stood out and contained a known pigmentation gene, *TYRP1* (Table S2). To test if additional loci beyond *TYRP1* influenced quantitative color phenotypes in black bears, we reran the analysis using the R153C genotype from the qPCR (see below) as a third covariate.

We estimated the length of the haplotype containing the derived allele within the Nevada population using HAPLOVIEW [14] using the default algorithm for estimating linkage disequilibrium (LD) [36]. We then viewed this region of the genome for all samples using HAPLOSTRIPS [13]. The Nevada population was selected based on the *runtc* (see below and Figure S5) analyses which indicated it was geographically near the source population for the R153C *TYRP1* population.

### Candidate Gene Analysis

To evaluate deleterious protein coding variation in bears, we downloaded coding sequences (CDS) for 13 candidate coat color genes (Table S3) from the dog genome *(Canis familiaris*; CanFam3.1; Table S2). Exons were queried against genome assemblies of *U. americanus, U. arctos*, and *U. maritimus* [10, 37, 38] using the blastn and dc-megablast versions of NCBI BLAST+ [39, 40]. We extracted the resulting sequences from the phased genome assemblies using SAMTOOLS v1.9 [41] and BCFTOOLS and concatenated exons to form haplotypes. We translated each haplotype and identified missense and nonsense polymorphisms within *U. americanus*, between *U. americanus* and *U. arctos*, and between *U. arctos* and *U. maritimus*. We tested each variant for deleterious effects using PROVEAN, PolyPhen2, and SIFT [42-44]. For any variant where two of the programs predicted a deleterious variant, we identified the individuals and coat color phenotypes of those animals to assess if there was concordance between *U. americanus* animals with brown coats and the alternate allele. Only the previously identified *TYRP1*^*R153C*^ showed such a concordance. We constructed a neighbor joining tree from the haplotypes (with introns) using GENEIOUSPRIME [45] to visualize the gene tree.

### Genotyping TYRP1 R153C

We genotyped 906 *U. americanus* samples for the *TYRP1*^R153C^ variant using a qPCR assay.

The primers included (5’ to 3’):

F: CCTTGAAGTCAGGAGAAACC

R: CTGGTCGCAATGACAAAC

The probes included (5’ to 3’):

reference allele probe: FAM-ATGGCGAAGCGCACAATTC-BGQ1

alternate allele probe: TET-ATGGCGAAGTGCACAAT-BHQ1

Primers and probes were ordered from IDT. Reaction conditions had final primer and probe concentrations of 1µM and 0.2 µM, respectively. The thermocycler conditions in a BioRAD CFX96 included: 95°C for 3min, and 40 cycles of 95°C for 15sec then 57.9°C for 30sec. For samples where both hair color phenotypes and R153C genotypes were available (n = 317), we tested if reflectance was significantly different between genotypes using two-sided t-tests in R v3.6.0 [46].

### Functional Validation of TYRP1 Alleles

To test if the *TYRP1* variants we identified in American black (R153C) and brown (R114C) bears caused a change in pigmentation phenotype, we first used site-specific mutagenesis to generate recombinant retroviruses to co-express wildtype (WT) or variant versions of human and mouse TYRP1 with a drug resistance marker for stable cell line selection. Specifically, we first subcloned a WT *TYRP1* cDNA based on the human sequence (pCDNA3-TRP1, a gift to MS Marks from W Storkus, University of Pittsburgh, Pittsburgh, PA, USA) into the pBMN-I-Hygro retroviral vector [47]. The R153C and R114C variants were generated individually by overlapping PCR using synthetic oligo nucleotides bearing the desired mutations (denoted by asterisks in the sequences below).

The oligos for R153C were (5’ to 3’):

F: GGGCCCTGGATATGGCAAAGT^*^GCACAACTCACCCTTTATTTGT

R: ACAAATAAAGGGTGAGTTGTGCA^*^CTTTGCCATATCCAGGGCCC

The oligos for R114C were (5’ to 3’):

F: GACACAACTGTGGGACGTGCT^*^GTCCTGGCTGGAGAGGAGCTGC

R: GCAGCTCCTCTCCAGCCAGGACA^*^GCACGTCCCACAGTTGTGTC

The flanking oligos for both products were (5’ to 3’):

F: CCTCTAGACTGCCGGATCCATTTAAATTCGAATTCCTGCAGG

R: GGAATCAAAGTTGCTTCTGGATCCCATCAAGTCATCCGTGCAG

The R153C mutation was generated by combining a 707bp and a 386bp product to obtain a 1,050bp product. The R114C mutation was generated by combining a 590bp and a 503bp product to obtain a 1,050bp product. The PCR products bearing the mutations were cloned individually as 1,013bp BamHI fragments into the BamHI site in the TYRP1pBMN-I-Hygro construct after excising the WT BamHI fragment. The clones bearing the correctly oriented fragments were selected by HindIII restriction and confirmed by Sanger sequencing the BamHI junctions. The generated mutations were confirmed by Sanger sequencing.

The WT mouse *TYRP1* cDNA sequence (NCBI accession NM_031202.3) was also used to generate the mutant forms. The three versions (WT, R153C, and R114C) were synthesized individually by Synbio Technologies as XhoI/HpaI fragments (of 2270bp) and cloned into the XhoI/SnaBI site of the pBMN-I-Hygro vector. The three sequences were confirmed by Sanger sequencing.

Immortalized melan-b melanocytes derived from mice homozygous for the *TYRP1*^*b*^ allele were cultured as described [48, 49]. Melan-b cells were infected with recombinant retroviruses encoding *TYRP1* variants (described above). Briefly, Plat-E cells [50] were cultured and transfected with the retroviral vectors described above, empty pBMN-I-hygro, or pBMN-I-hygro expressing mCherry-tagged human STX13 [51] using Lipofectamine 2000. The medium was replaced with fresh melanocyte medium the following day and cell supernatants were harvested 48-72 h post-transfection as described [52]. The medium, containing recombinant retroviruses, was used directly to infect melan-b cells as described [53] except that cells were seeded the previous day in a 10-cm dish. Two days later the medium was supplemented with 500 mg ml^-1^ hygromycin B to select for infected cells. Antibiotic-resistant cells were used directly as polyclonal cell populations for experiments. Two separate stable lines per construct were selected and included in the analyses.

We performed spectrophotometric assays or melanin and protein content as described [53, 54]. Briefly, melanocytes in a confluent 10-cm dish were harvested by trypsinization. Cells were counted, and duplicate samples of 10^6^ cells for each experimental condition per experiment were washed once with RPMI + 10% FBS and twice with ice-cold PBS, and then resuspended in 50 mM Tris-HCl, pH 7.4, 2 mM EDTA, 150 mM NaCl, 1 mM diothiothreitol, and cOmplete Protease Inhibitor Cocktail Tablets (Roche). Cells were probe-sonicated on ice using the Sonic Dismembrator Model 100 (Fisher Scientific), and sonicates were fractionated into supernatant and pellet fractions by centrifugation at 20,000 x g for 15 min at 4°C. Protein concentrations in supernatants were estimated using the DC colorimetric protein assay (Bio-Rad Laboratories) with a standard curve derived using differing concentrations of BSA. Melanin-containing pellets were washed in 500 μl 50% ethanol, 50% ether, collected by centrifugation at 20,000 x g, and resuspended in 1 ml of 2 M NaOH, 20% DMSO. Melanin was solubilized by heating at 60°C for 40–60 min and quantified using spectrophotometry to measure Optical Density (OD) at 492 nm. OD values were normalized to total protein content in each lysate.

We performed imaging and analysis as described [55]. Briefly, melanocytes were seeded onto Matrigel-(Thermo Fisher) coated coverslips 24–48 h before fixation. Cells were fixed with 2% formaldehyde in PBS for 20 min, washed briefly with PBS, then labeled with TA99/Mel-5 antibody to TYRP1 (American Type Culture Collection, Rockville, MD) diluted in blocking buffer (PBS, 0.1% BSA, 0.02% saponin) for 1 h at room temperature (RT). After a 15-min wash in PBS, cells were incubated with Alexafluor 488-labeled donkey anti-mouse IgG (Jackson Immunoresearch) diluted in blocking buffer for 30 min at RT. Samples were washed for 15 min in PBS, mounted with Prolong Gold Antifade Mountant (Invitrogen), and analyzed by epifluorescence microscopy on a DMI 6000B microscope (Leica Biosystems) equipped with a 63x plan apochromat (Plan Apo) objective (1.4NA) and a Hamamatsu Photonics ORCA-Flash4.0 sCMOS digital camera. Both fluorescence and bright field images were acquired as a z-series with 0.19-μm steps and were deconvolved using the blind deconvolution algorithm of Microvolution software (Bio-Vision Technologies) and ImageJ. The area of overlap between TYRP1 and pigmented melanosomes in the periphery of melanocytes was quantified using a threshholding approach in ImageJ on deconvolved fluorescence images as detailed. Final images were composed using Adobe Photoshop software [53].

Statistical analyses including two-way ANOVA and Tukey’s test were performed using Graphpad Prism 9 for MacOS.

### Allele Age

We estimated the age of *TYRP1*^*R153C*^ using *runtc* [23], a program that estimates the time of first coalescence to the nearest variant on a chromosome. To increase sample size for this analysis, we used the combined high and imputed low coverage data for HiC_scaffold_24. We reran the GLIMPSE pipeline on the low coverage samples where the high coverage reference panel was not first filtered for MAF, thereby leaving more singletons in the reference dataset for coalescence with the focal allele. We estimated the coalescence time for all variants on the scaffold with all range wide samples together, as well as separately in the western lineage, SEAK, and eastern populations separately, using a mutation rate of 1×10^−8^ and the recombination rate from the dog genome, 0.97cM Mb^-1^.

### Demography

We inferred the demographic history of *U. americanus*, then tested if spatial patterns of *TYRP1*^*R153C*^ alleles were due to neutral or adaptive processes given the demography. We first assessed *U. americanus* population structure using PCA. We aligned a previously generated RAD-Seq dataset [7] to the *U. americanus* reference genome and called SNPs using STACKS v2.53 [56], including filters for one SNP per RAD-tag, >80% genotyping success per locus, and MAF ≥ 0.05; this resulted in 37,422 SNPs. We pulled down those loci from the high coverage WGS samples to increase the sample size (n = 117). We ran two PCA analyses in PLINK using the RAD-seq+hcWGS and imputed lcWGS datasets, and observed concordant patterns (Figure S12) between datasets and with previous analyses [7].

We estimated change in effective population size (N_e_) through time using MSMC2 [57]. For this analysis we used scaffolds 2–37, and made individual mask files with the bamCaller.py script, then used the phased datasets to make genome-wide masks with SNPable [58]. We generated input scripts using generate_multihetsep.py. Output was converted to years and number of individuals using a mutation rate of 1×10^−8^ [59] and a generation time of 6.5 years [60]. We ran MSMC on three samples (six haplotypes) from both the eastern and western lineages, and the admixed SEAK population, and PSMC’ on each sample (two haplotypes). Historic N_e_ was approximately 30,000 (Figure S14). From 100–10kya N_e_ declined to 3,000 in the western lineage, yet remained constant in the eastern lineage. N_e_ also declined in Alaska from 50–10kya to approximately 8,000. MSMC plots suggest N_e_ increased since the Last Glacial Maximum (18– 22kya). This demographic history provides an explanation for the low haplotypic diversity in Nevada as the ancestral and causative alleles sit on highly homogeneous haplotypes (Figures S3, S8, S10). Using the output from MSMC, we further estimated the cross-coalescent model and input results into MSMC-IM [24] to estimate the patterns and timing of population divergence (using MSMC estimates), and rate of gene flow over time (using the PSMC’ estimates; Figure S13). Estimates may be inaccurate in the most recent time lags of MSMC analyses warranting caution when estimating contemporary N_e_. Our data appears to have a limit for inferences younger than 1–5kya.

While our demographic model infers broad patterns of lineage relatedness, it does not identify the directionality of range expansion; thus, we used the RAD-seq+hcWGS dataset to understand patterns of range expansion via the directionality index (Ψ) [61]. We removed geographic sites with fewer than three samples for a total of 111 samples across 23 populations. We input genotypes and geographic data into the *rangeExpansion* package in R, then calculated Ψ for all population pairs using the get.all.psi function. We determined the significance of each pairwise Ψ estimate by calculating the standard error of the upper triangle of the matrix excluding the diagonal, then calculating a Z-score for each pairwise comparison. To visualize the range expansion patterns, we plotted pairwise values where Z ≥ |5| within the western and eastern lineages, and between the two lineages and SEAK.

### Selection Tests

We calculated pi (π) using VCFTOOLS [62] in 1kb sliding windows with a 100bp step size from the imputed low coverage WGS data using samples from the Nevada (western lineage; n = 38), SEAK (n = 95), and Appalachian Mountains (representative of eastern lineage; n = 38) populations. We further calculated *F*_ST_ between black and brown animals within either the Nevada or SEAK populations using the same sliding window scheme.

We tested for a selective sweep surrounding *TYRP1* using both integrated haplotype score (iHS) and extended haplotype homozygosity (EHH) [63] implemented in selscan v2 [64, 65]. The iHS analysis was run on scaffold 24 of the *U. americanus* reference genome within the Nevada population, then normalized based on allele frequency using the norm function within selscan 2. No sites within the haplotype block carrying *TYRP1*^*R153C*^ had iHS scores greater than |2|. We ran the EHH analysis in both the Nevada and SEAK populations using the --keep-low-freq flag.

Further, we identified 15 sites on each of the 36 long scaffolds (or 540 sites each) that matched the derived allele frequency of R153C within each test population and ran EHH on each of those sites. We compared the decay curves of R153C to these allele frequency matched background loci as a test if the decay in haplotype homozygosity was within the range of the background.

While decay curves between the ancestral and derived alleles suggested extended haplotype homozygosity for the derived R153C, the background matched samples showed the overall scores were not out of range for the background sites. Both the low N_e_ within the Nevada population and low selection coefficient complicate the analysis of selection, as simulation studies suggest low power to detect selection under these conditions [66]. Moreover, we chose not to use cross-population sweep tests due to the decrease in power with increasing divergence, and our estimates that the eastern and western lineage populations diverged ∼15.8k generations ago [64].

We further tested for possible drivers of selection using BayEnv2 [27, 67]. Using the combined RAD-Seq, hcWGS, and lcWGS data at the RAD loci, we clustered the data into 24 geographic areas, identified a centroid longitude and latitude coordinate, then extracted values of 11 climate variables and altitude for each coordinate. The climate variables from the CliMond dataset [68, 69] included: bio1 (annual mean temperature), bio3 (isothermality), bio4 (temperature seasonality), bio5 (max temperature of warmest week), bio6 (min temperature of coldest week), bio10 (mean temperature of warmest quarter), bio11 (mean temperature of coldest quarter), bio12 (annual precipitation), bio19 (precipitation of coldest quarter), and bio20 (annual mean radiation). Beyond the climate variables, altitude, latitude, and longitude were all tested as selective forces. Finally, we used the presence or absence of *U. arctos* on the landscape as a factor. For each geographic location, we scored the site with the presence of *U. arctos* before range contraction due to human activity based on maps of the expected species distribution [70]. We prepared the covariance matrix using PGDSpider [71], and the environmental matrix according to the BayEnv2 manual. We ran 100,000 iterations of the model and analyzed the Bayes Factors for the alternative hypothesis that the variable of interest was a selective force on allele frequency of *TYRP1*^*R153C*^.

### Gene Flow Simulation

To test if a model of gene flow without selection could produce the spatial gradient of declining frequency of the alternative allele along a south-to-north axis is the western range, we used the forward genetic simulation program SLiM3 [72]. We modeled four populations, broadly representative of Nevada, Arizona/New Mexico, Idaho, and SEAK. These population were selected as we have at least one high coverage genome from each population (Table S1). We reran MSMC-IM on one genome from each population for all pairwise combinations to estimate historic gene flow rates to use as input in our model (Figure S12).

Our model seeds a new variant (i.e., R153C) at a frequency of 1% into the Nevada population at generation 0, then tracks allele frequency within each population for 1,440 generations such that we get a simulated estimate to compare to observed data. Our data suggests that R153C has a dominance coefficient (*h*) somewhere greater than 0.5, but not 1; thus, we varied *h* from 0 to 1 using 0.1 increments. Further, we ran each model with a selection coefficient (*s*) of either 0, 0.005, or 0.01. MSMC-IM estimates bidirectional gene flow, thus for each population pair we use the peak estimated rate (Figure S12) since the LGM. We ran three different models varying the selection coefficients (*s* = 0, 0.005, or 0.01). From generations 980–1,310, population size increased in each population at a per generation rate of 10^−6^ which was a rate approximated from our MSMC results. At generation 1,390 (i.e., 50 generations prior to the end of the simulation and meant to represent anthropogenic influences on bear populations), we imposed population specific bottlenecks given estimates from the literature. Specifically, the Nevada population was reduced to 100 [73], and the Arizona/New Mexico, Idaho, and SEAK populations were reduced to 5,000, 7,000, and 4,000 respectively [7]. We ran 1,000 iterations of each model. We recorded the number of simulations in which the allele went extinct in all populations or was still polymorphic following 1,440 generations (no simulations observed the derived allele fixing in all populations). For simulations where the locus remained polymorphic, we averaged the frequency within each population across simulations, and compared to the observed frequency from our qPCR assay and contour plots of percent of black animals in the population [5].

This simulation with 33 different scenarios (*h* by *s* combinations) identified that both increasing values of *h* or *s* resulted in increased allele frequencies and decreased probability of allele extirpation from the landscape (Figure S7). We reran the simulation with an *h* of 0.75 for the final model (Figure 6).

## ACKNOWLEDGEMENTS

We thank Omar Skalli for microscopy training and Jay Puckett for assistance with maps. AP was supported by NIH R35 GM134957-01 and American Diabetes Association Pathway to Stop Diabetes grant 1-19-VSN-02. Any use of trade, firm, or product names is for descriptive purposes only and does not imply endorsement by the U.S. Government.

## DATA ACCESSIBILITY

WGS data will be deposited to the NCBI SRA upon acceptance.

