## Supplemental Figures for "Genetic architecture and evolution of color variation in American black bears"

**SUPPLEMENTAL TABLES**

**Table S1-** Samples used for high (hcWGS) and low (lcWGS) coverage whole genome sequencing with SRA accessions.

**Table S2-** All GWAS hits for brown color in *U. americanus* with $P < 10^{-8}$ and genes under the peak.

**Table S3-** Genomic positions for 13 candidate genes associated with hair pigmentation. The scaffold, and start and stop position for each exon and the full gene are listed for *U. americanus*, *U. arctos*, and *U. maritimus*. Note that the start and stop positions are oriented correctly if the gene is in the reverse orientation.

**Table S4-** Missense and nonsense amino acid substitutions identified within *U. americanus* WGS for 13 candidate genes. For each variant the Provean, PolyPhen2, and SIFT scores and associated predictions are listed.

**Table S5-** Missense and nonsense amino acid substitutions identified in eight of 13 candidate genes between *U. americanus* and *U. arctos*. For each variant the Provean, PolyPhen2, and SIFT scores and associated predictions are listed.

**Table S6-** Missense and nonsense amino acid substitutions identified in six of 13 candidate genes between *U. arctos* and *U. maritimus*. For each variant the Provean, PolyPhen2, and SIFT scores and associated predictions are listed.

**Table S7-** Bayes Factors from BayEnv2 testing 15 variables as the potential selective force for the frequency of *TYRP1* R153C among populations across the range of American black bears.

**SUPPLEMENTAL FIGURES**

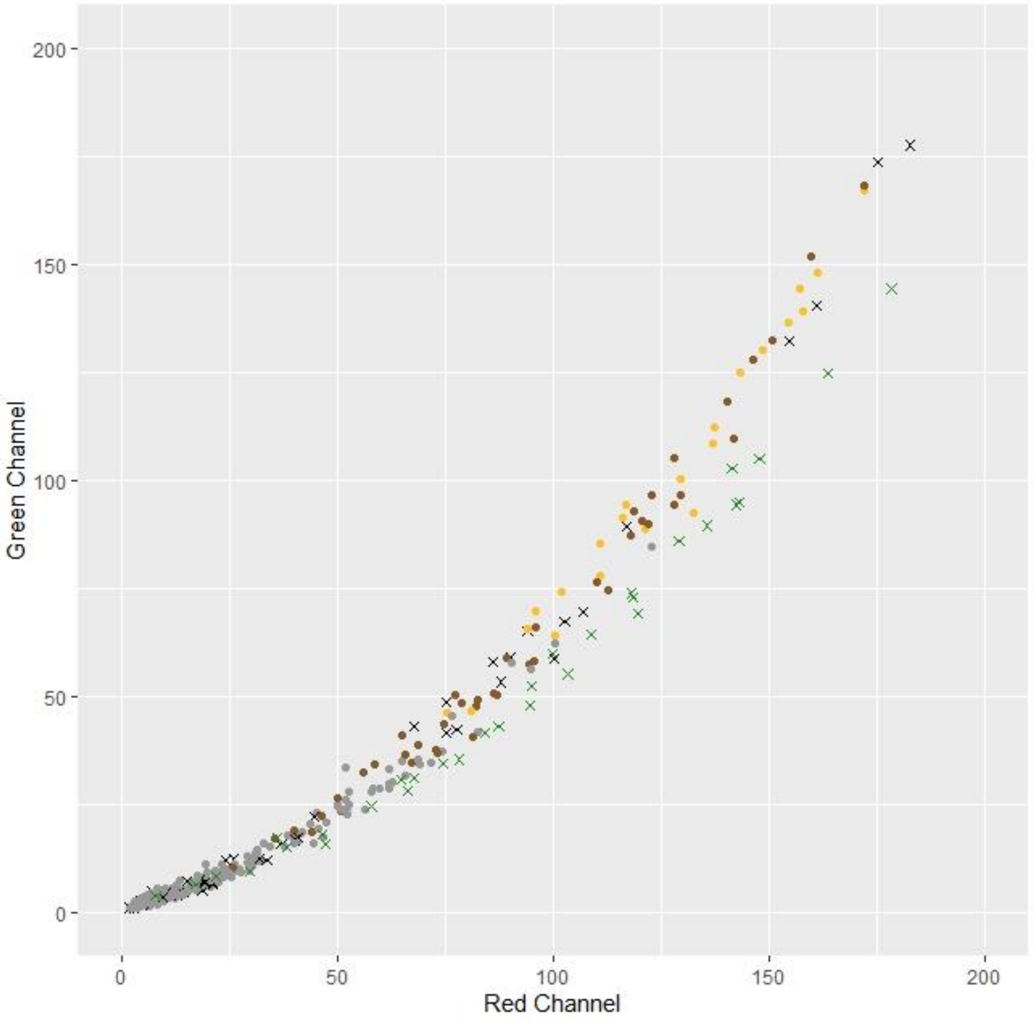


**Figure S1-** Hair reflectance scores for the red (x-axis; range 0–255) and green (y-axis; range 0–255) channels for *U. americanus* and *U. arctos*. Sample data for *U. americanus* is colored by *TYRP1* R153C genotype, including: homozygous ancestral (GG; grey), heterozygous (GA; brown), homozygous derived (AA; gold), and no genotype data (black "X"); whereas, all *U. arctos* samples are shown as green "X."

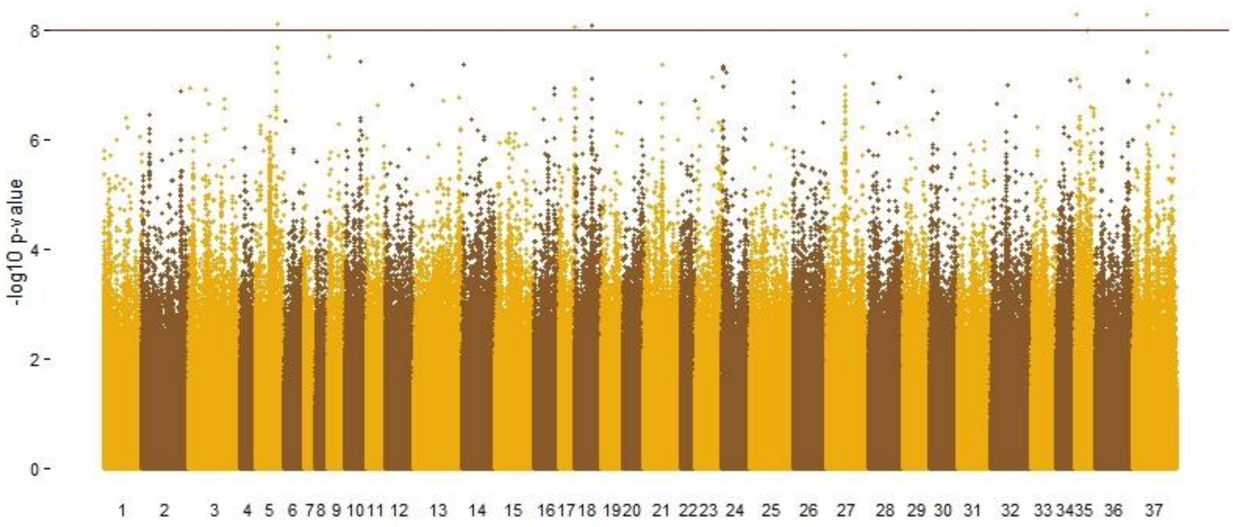


**Figure S2-** Manhattan plot of *U. americanus* WGS data with *TYRP1* R153C causative SNP serving as a covariate when comparing hair reflectance to genotype. A genome-wide significance cutoff value of $10^{-8}$ was selected (brown horizontal line).

[See separate pdf]

**Figure S3-** *Ursus americanus* haplotypes from phased high and low coverage WGS (see Table S1) on scaffold 24 encompassing the 97kb haplotype identified in the Nevada population containing *TYRP1*. Alternate alleles shown as black lines for each sample (represented as a row). Relationships among haplotypes represented as a gene tree on far left, geographic location of samples noted by leftmost colored bar (western lineage: orange; Southeast Alaska: purple; eastern lineage: blue), and qPCR genotype of R153C represented by rightmost colored bar (homozygous ancestral: black; heterozygous: brown; homozygous derived: gold).

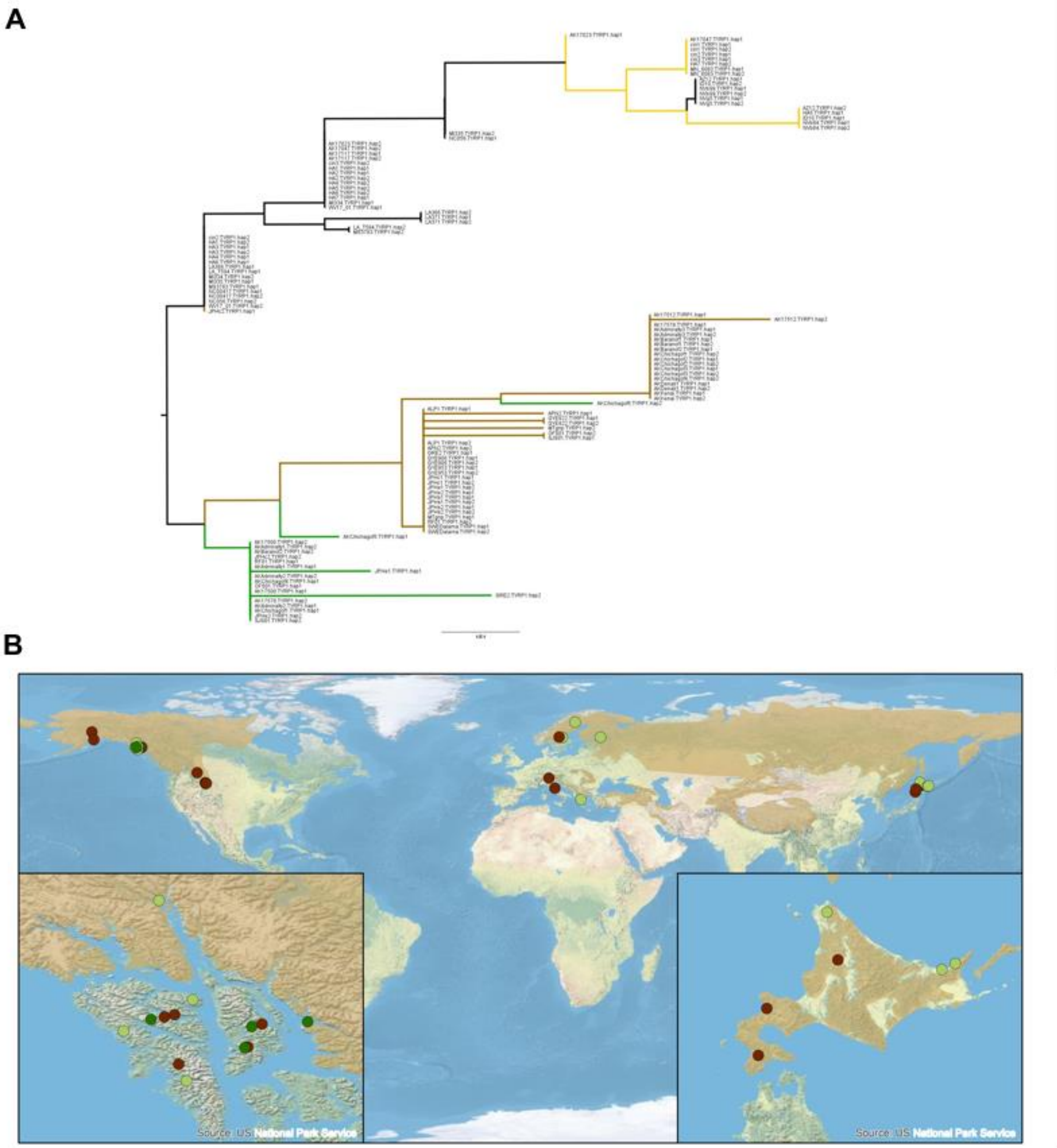


**Figure S4-** (A) Unrooted neighbor joining haplotype tree of *U. americanus* and *U. arctos TYRP1* haplotypes (including exons and introns). Haplotypes carrying causative variants within *TYRP1* are denoted by color with *U. americanus* (American black bear) in black (R153 ancestral allele) or gold (C153 derived allele), and *U. arctos* in brown (R114 ancestral allele) or green (C114 derived allele). (B) Map of the global distribution of R114C in *U. arctos* individuals sequenced to high coverage (see Table S1), where brown circles indicate homozygous ancestral, light green are heterozygous individuals, and dark green are homozygous derived. Left inset map highlights Southeast Alaska, USA; right inset map highlights Hokkaido, Japan.

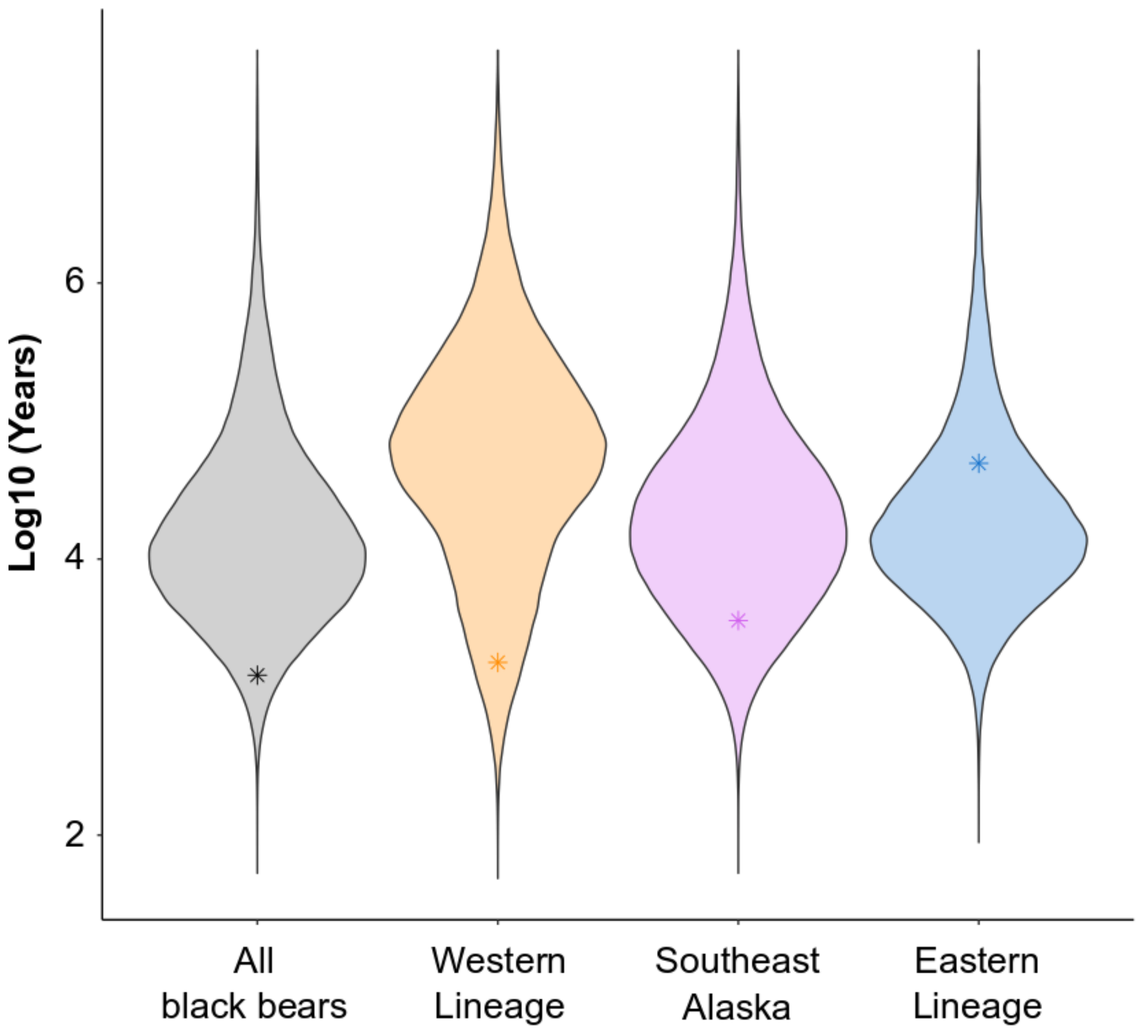


**Figure S5-** Coalescence time estimates from all variants on *U. americanus* scaffold 24 for all samples (grey; n = 185), western lineage (overrepresented by the Nevada population; orange; n = 44), Southeast Alaska population (purple; n = 84), and eastern lineage (blue; n = 57). The star (*) within each violin plot represents the estimate of the *TYRP1* R153C causative variant.

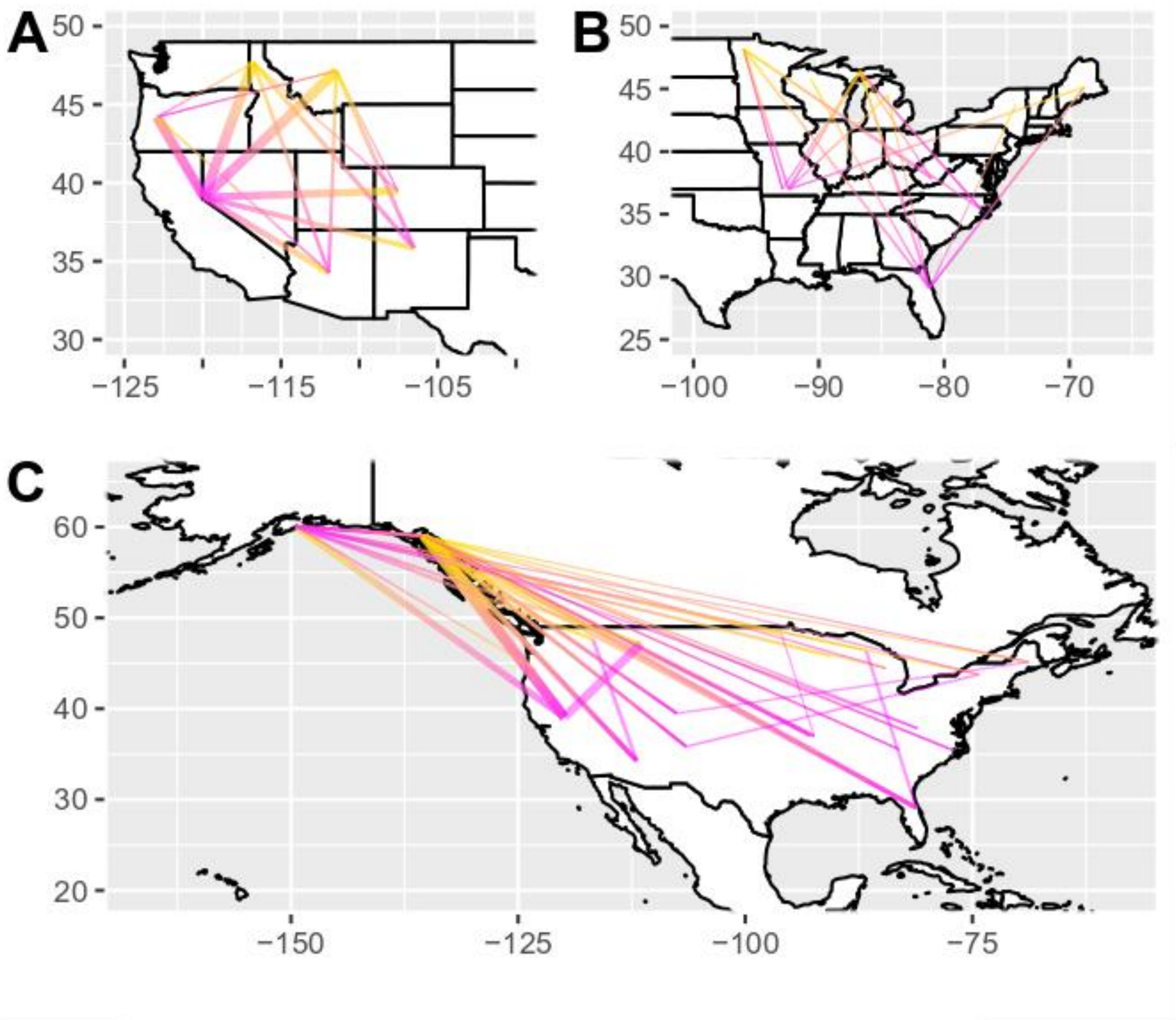


**Figure S6-** Estimates of regional *Ursus americanus* range expansions based on pairwise ψ statistics (A) within the western lineage populations, (B) within the eastern lineage populations, and (C) among the western and eastern lineages and Alaskan (both Southeast and Kenai) populations. Lines show directionality from inferred source (magenta) to sink (gold) populations, where thickness was scaled to the *Z*-score when the absolute value was greater than five.

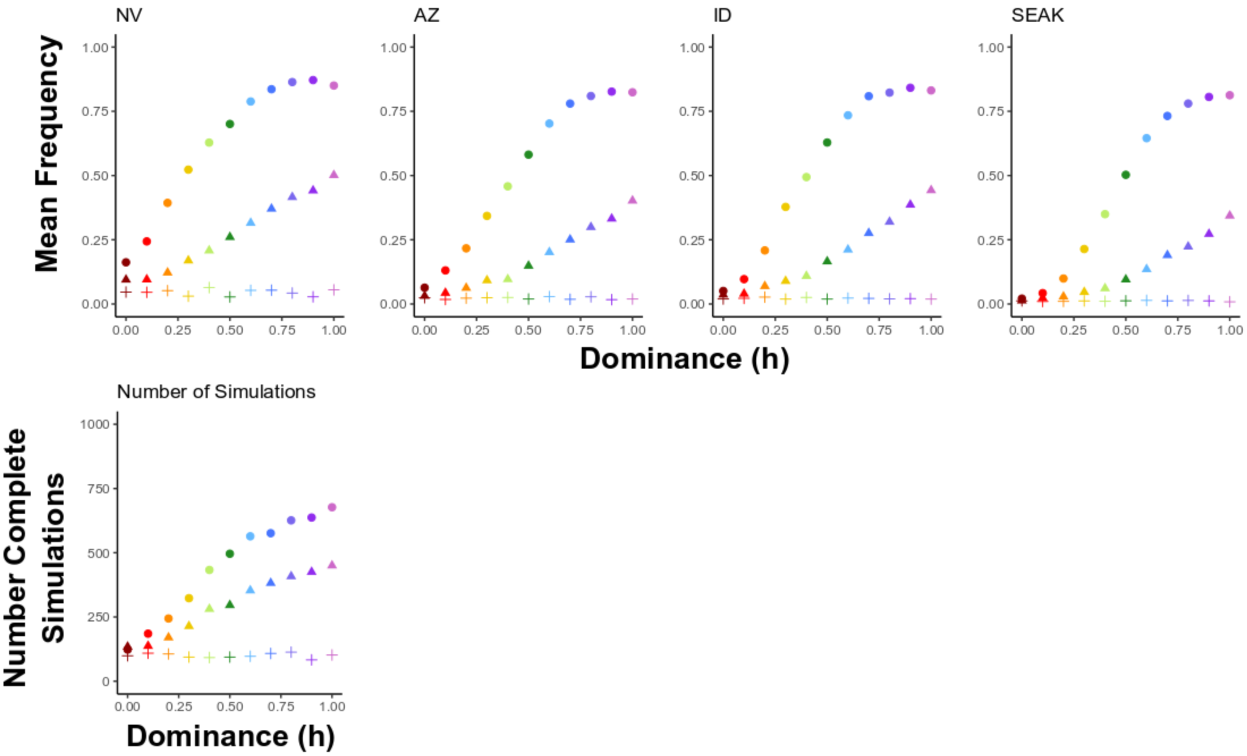


**Figure S7-** (Top Row) SLiM3 simulations of the final allele frequency of the derived allele (R153C) under simulations where the dominance (*h* varying from 0 to 1 in 0.1 increments; denoted by colors) and selection (*s*; denoted by symbols: 0- plus sign, 0.005- triangle, 0.01- circle) coefficients varied. Effective population size, growth rate, and migration were constant between simulations. (Bottom Row) The number of simulations (out of 1,000) for each combination of *h* and *s* (33 scenarios) where the derived allele did not go extinct. Only population frequency values from these simulations were averaged to estimate data in the plots on the top row.

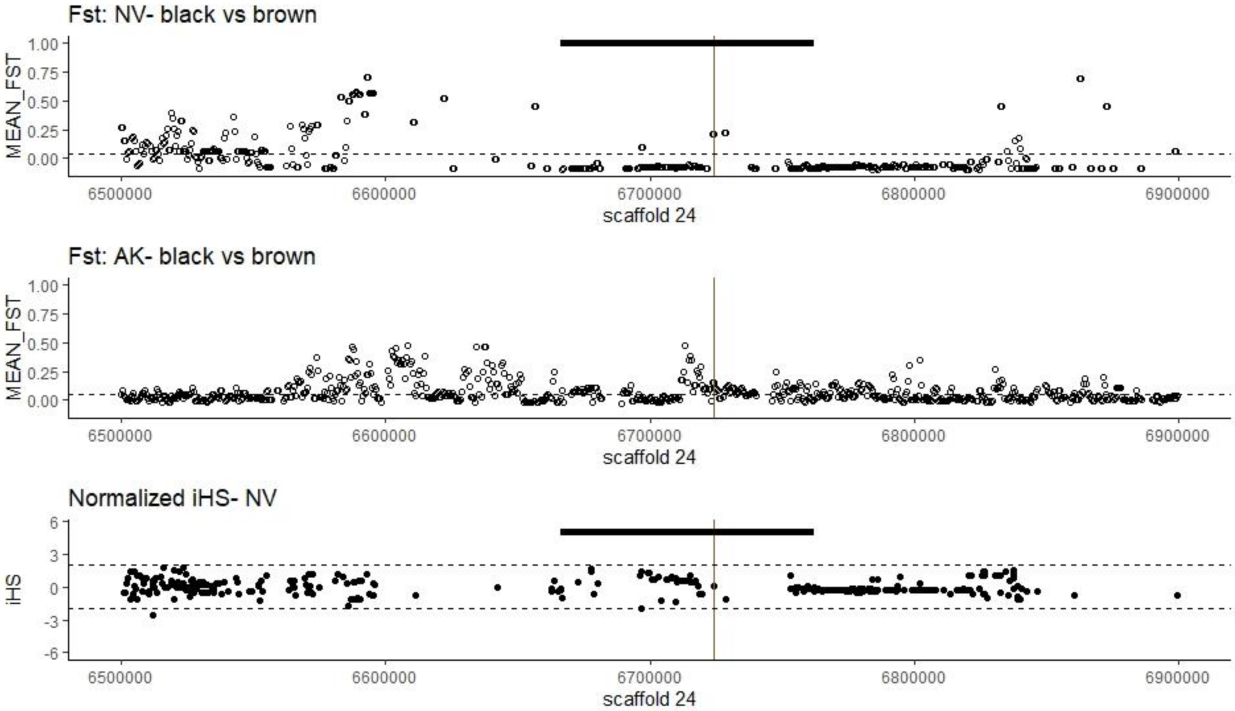


**Figure S8-** Genetic differentiation ($F_{ST}$) and selection (integrated haplotype score, iHS) tests across scaffold 24 of the *U. americanus* genome. (Top and middle panels) Pairwise $F_{ST}$ estimates in sliding windows (1,000bp window; 100bp slide) between black and brown coated animals within either the Nevada (top) or Southeast Alaskan (middle) populations, where average $F_{ST}$ for the scaffold is indicated by the dashed line. (Bottom panel) iHS was calculated in sliding windows (10k window, 2.5k slide) across scaffold 24 and normalized by allele frequency using selscan. The dashed lines represent iHS scores of |2| which is considered significant. In each panel, the vertical line represents the position of the R153C allele. The thick black bar in panels of the Nevada population show the length of the 97kb haplotype identified.

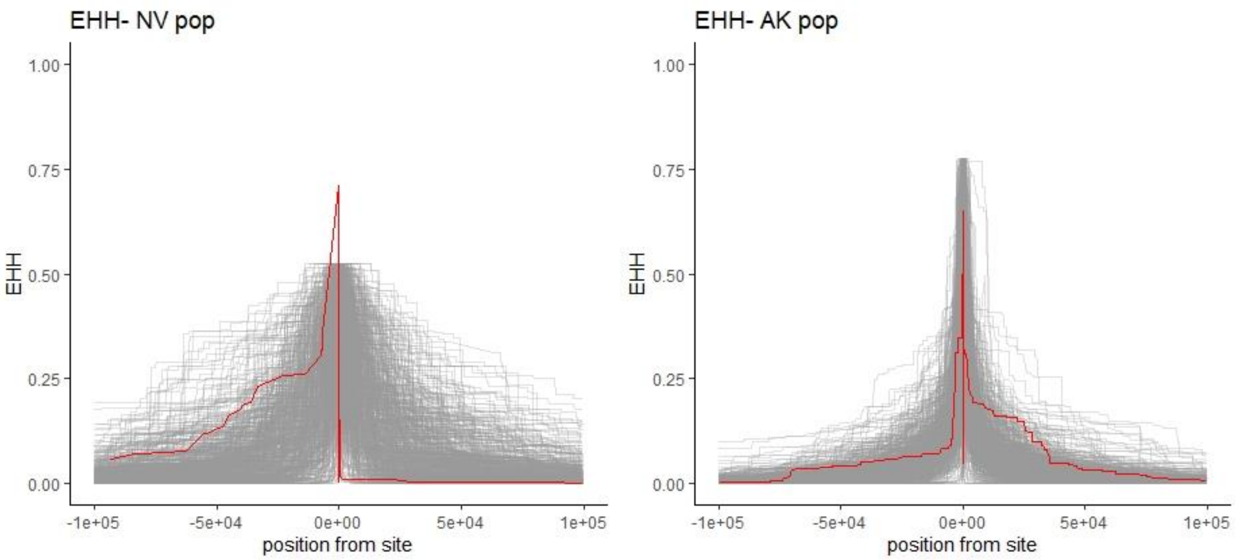


**Figure S9-** Extended haplotype homozygosity (EHH) of the causative *TYRP1*$^{R153C}$ allele (red line) in both the Nevada (left) and SEAK (right) populations compared to 540 allele frequency matched sites from throughout the genome of *U. americanus*.

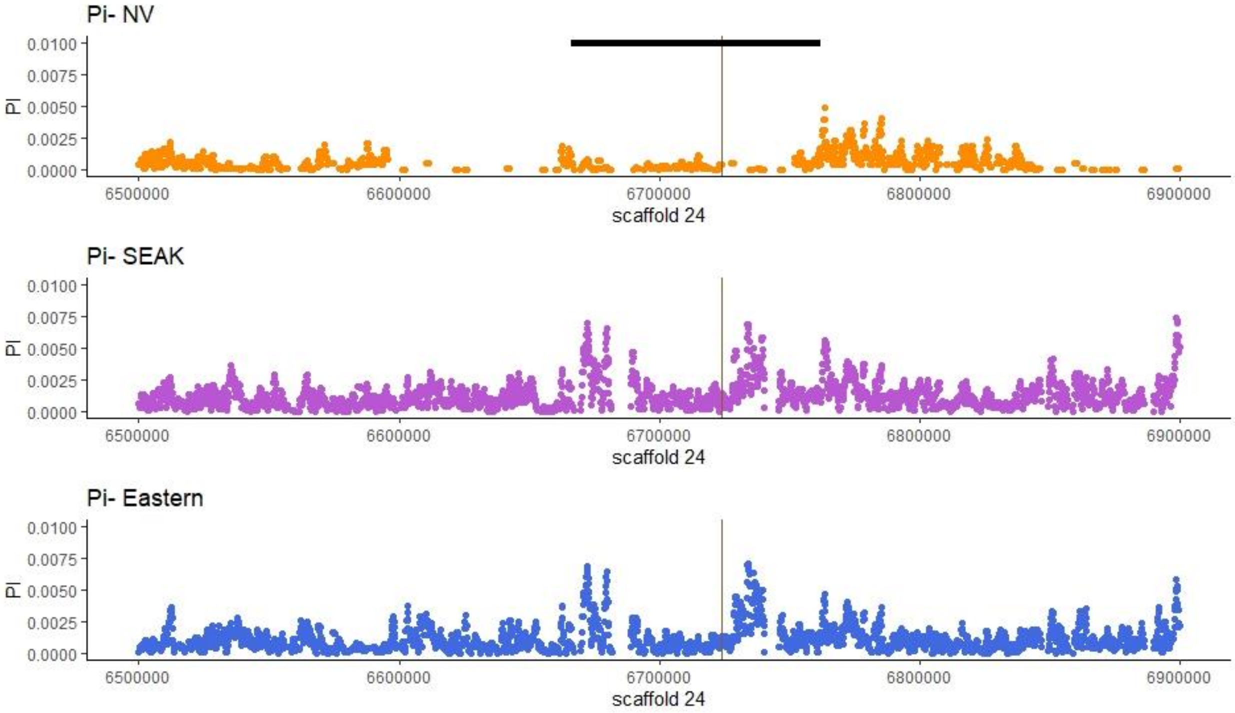


**Figure S10-** Nucleotide diversity within three different populations (top: Nevada in the western lineage; middle- Southeast Alaska; bottom- Appalachian Mountains in the eastern lineage) of *U. americanus* along scaffold 24. In each panel, the vertical line represents the position of the R153C allele. The thick black bar in the Nevada population shows the length of the 97kb haplotype identified; notably, diversity is low across the haplotype regardless of coat color phenotype.

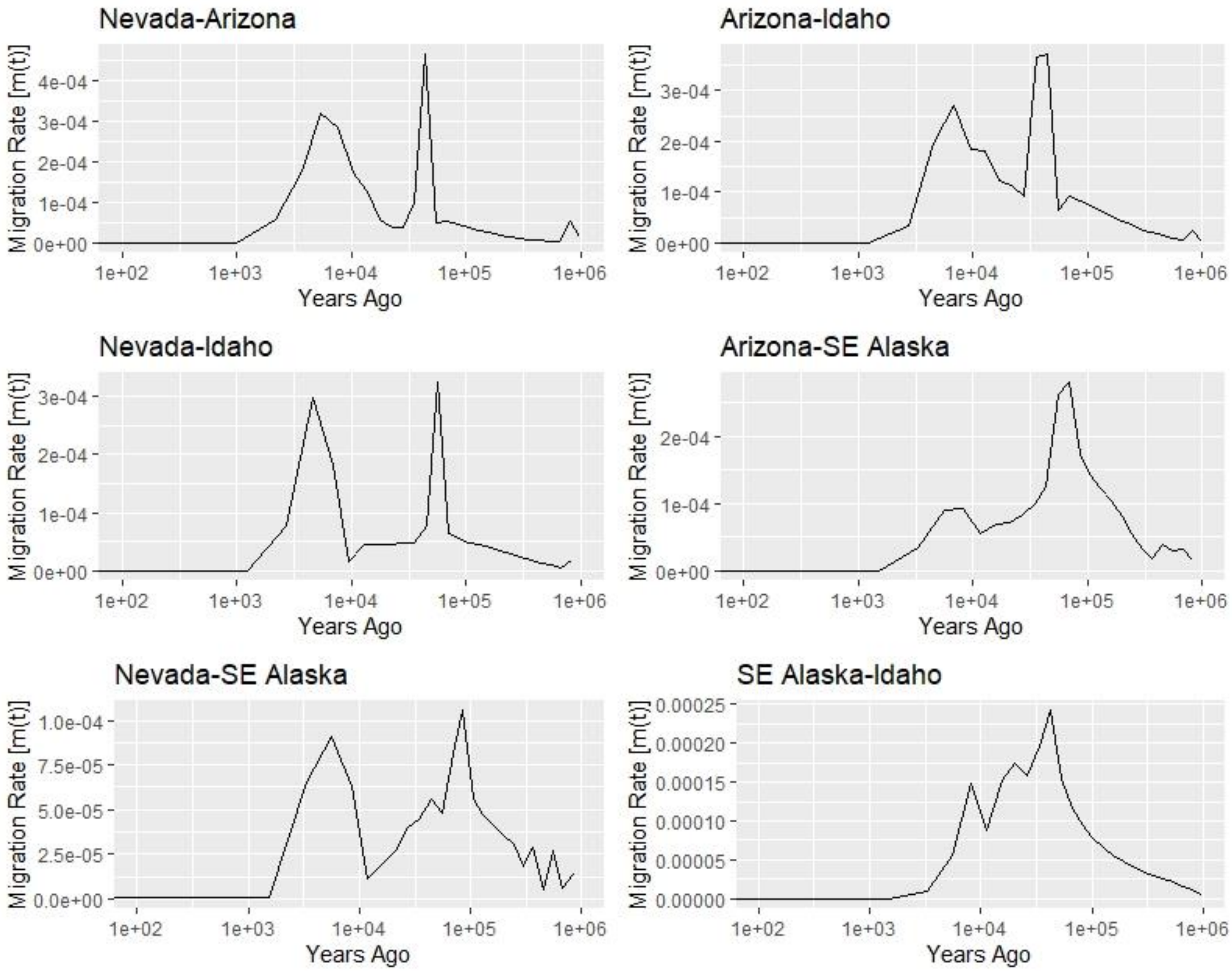


**Figure S11-** Estimated historic gene flow among four populations (Nevada, Arizona, Idaho, and Southeast Alaska) based on MSMC-IM estimates from a single WGS genome per population. The peak bidirectional migration between each population was used as the rate within the SLiM3 forward simulation models.

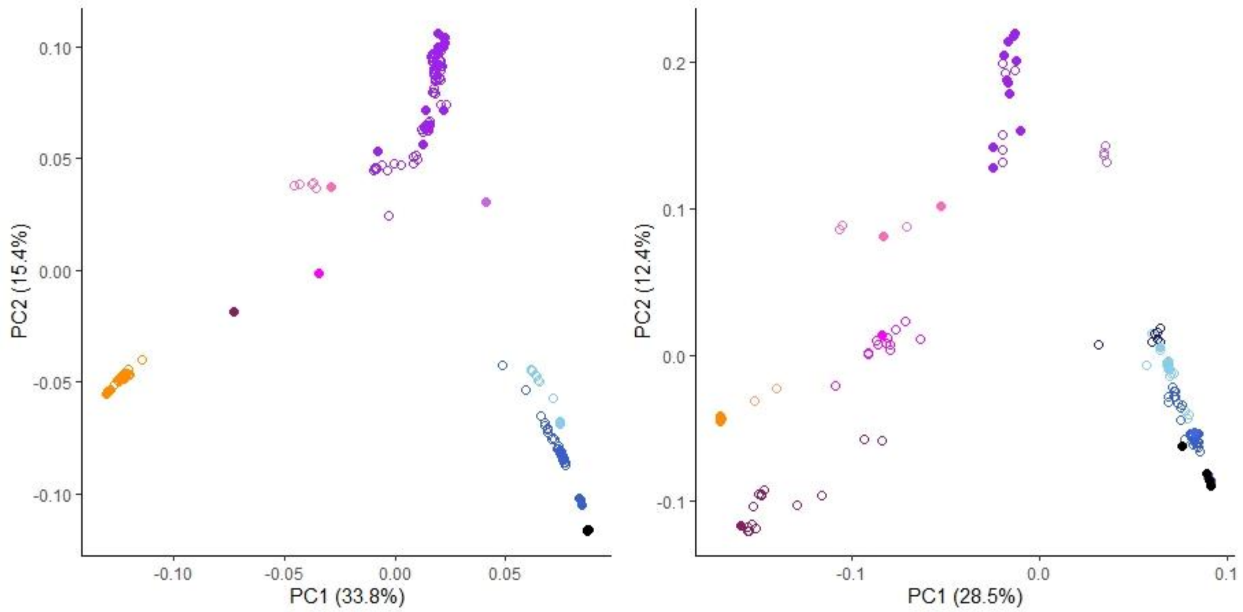


**Figure S12-** PCA of *U. americanus* genomic diversity across the range based on combined (left) high coverage WGS (closed circles) and imputed low coverage samples (open circles), and (right) 37k SNPs identified via RAD-seq. Sample colors correspond to: Nevada (orange), southern Rocky Mountains and southwest deserts (maroon), northern Rocky Mountains (magenta), southern SEAK (pink), SEAK (purple), Central Interior Highlands (navy), Great Lakes (light blue), southeast (medium blue), and Louisiana (black).

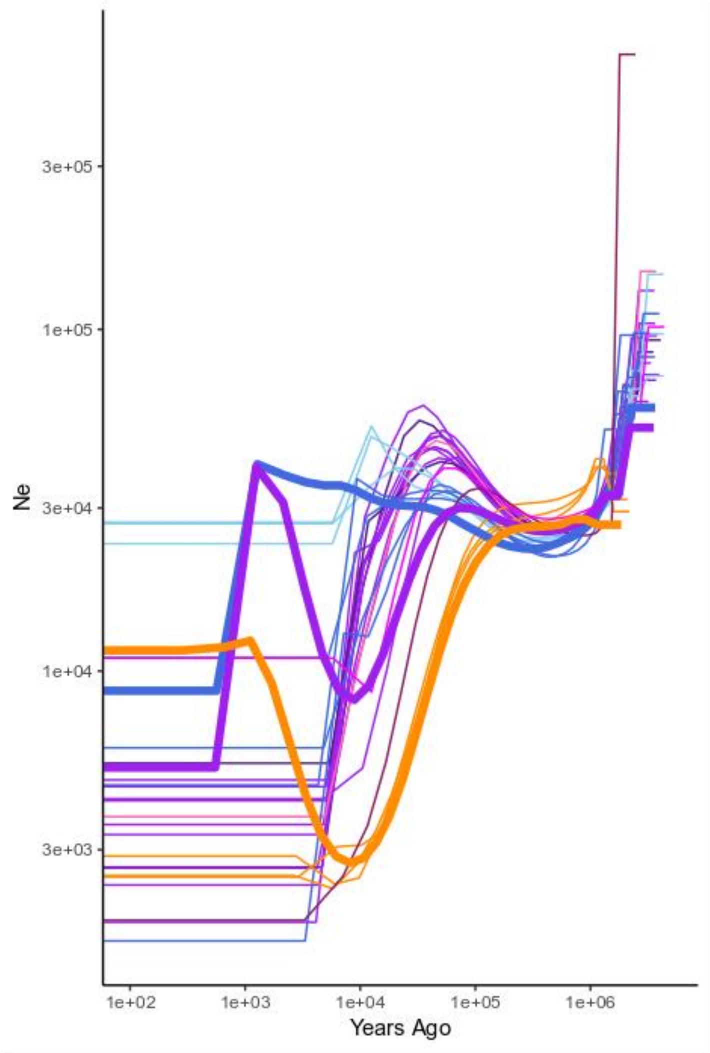


**Figure S13-** Change in effective population size ($N_e$) over time in *U. americanus*. Lineage estimates (western- orange; eastern- blue; Southeast Alaska admixed population- purple; thick lines) were estimated from six haplotypes; whereas, greater range variation was obtained from estimates from two haplotypes (thin lines). Colors include: Nevada (orange), southern Rocky Mountains and southwest deserts (maroon), northern Rocky Mountains (magenta), SEAK (purple), Great Lakes (light blue), and southeast (medium blue).
