## Supplementary figures and images for "Genetic architecture and evolution of color variation in American black bears"

### Figure S3

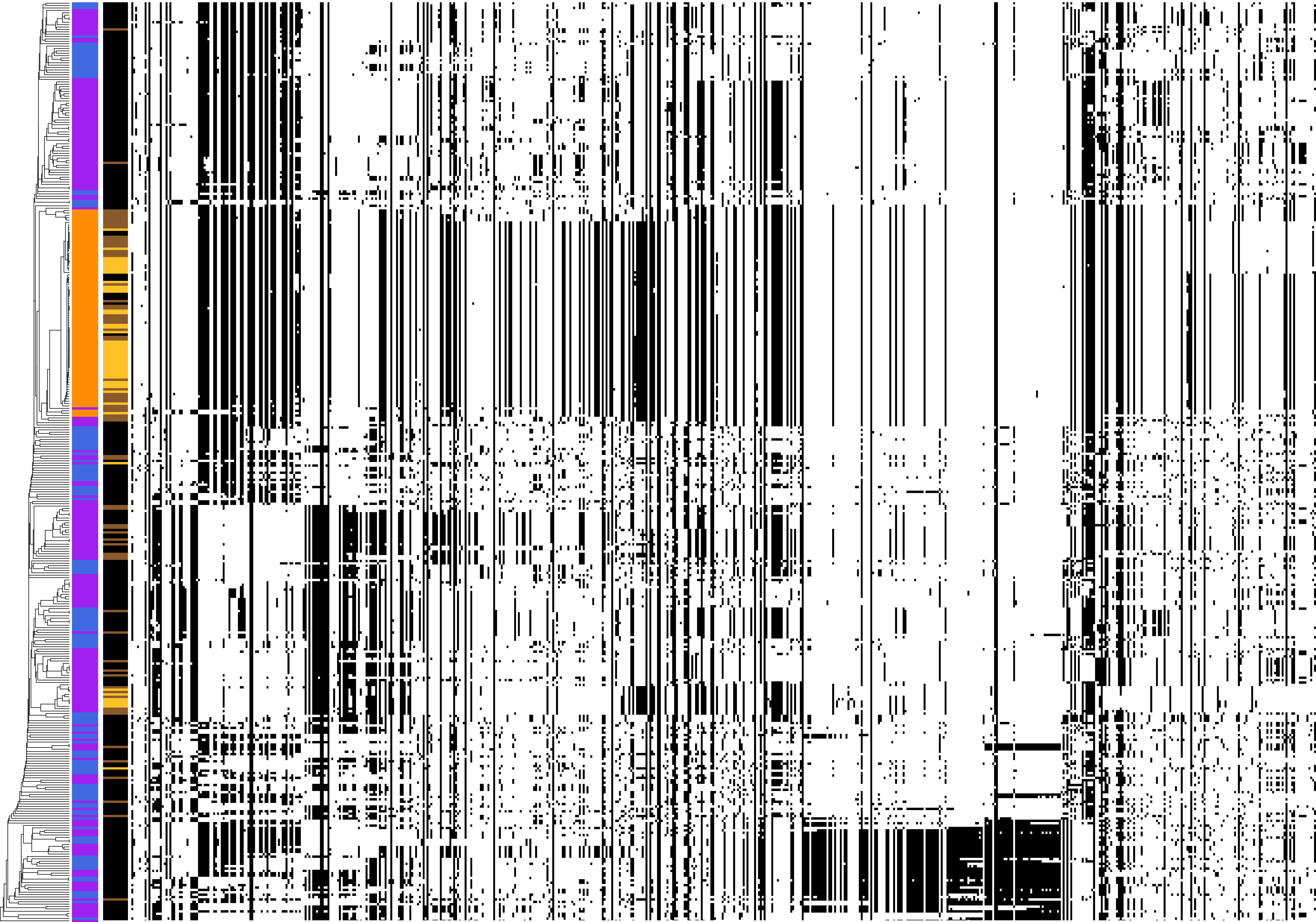
